# Multivalent interactions between eIF4G1, Pub1 and Pab1 drive the formation of protein condensates

**DOI:** 10.1101/2020.08.07.234443

**Authors:** Belén Chaves-Arquero, Santiago Martínez-Lumbreras, Nathalie Sibille, Sergio Camero, Pau Bernadó, Ma Ángeles Jiménez, Silvia Zorrilla, José Manuel Pérez-Cañadillas

## Abstract

Yeast eIF4G1 interacts with RNA binding proteins (RBPs) like Pab1 and Pub1 affecting its function in translation initiation and stress granules formation. We present an NMR and SAXS study of the intrinsically disordered region of eIF4G1, eIF4G1_1-249_, and its interactions with Pub1 and Pab1. The conformational ensemble of eIF4G1_1-249_ shows an α-helix within the BOX3 conserved element and a dynamic network of fuzzy π-π and π-cation interactions involving arginine and aromatic residues. The Pab1 RRM2 domain interacts with eIF4G1 BOX3, the canonical interaction site, but also with BOX2, a conserved element of unknown function to date. In contrast, the Pub1 RRM3 domain interacts with the RNA1-1 and BOX1 regions of eIF4G1. Mixtures of Pub1, Pab1 and eIF4G1 form micrometer-size protein condensates that require the presence of the eIF4G1 BOX1 element. These homotypic interactions suggest a double key mechanism of eIF4G1 regulation, important for understanding the architecture of stress granule cores.

## Introduction

Unicellular eukaryotes such as the yeast *Saccharomyces cerevisiae* have evolved mechanisms to regulate translation in response to different types of environmental stressors ^1^. The first reaction of yeast cells subjected to nutrient starvation, temperature, oxidative or chemical stresses is to globally shut-down protein synthesis by storing components of the translation machineries in membrane-less organelles known as stress granules (SG) ^2-5,6^. These cytoplasmic bodies are liquid-like condensates, contain proteins and RNA, and are generated by liquid-liquid phase separation (LLPS) ^7^. The importance of LLPS for biological systems is enormous and its discovery has given rise to parallel fields of research in chemistry, physics and biology ^7-9^. Many proteins present in SG contain intrinsically disordered regions (IDRs) and undergo LLPS when isolated at high concentrations ^10-12^. Moreover, the presence within the SG of proteins with both IDRs and folded domains involved in multivalent interactions is considered one of the key factors driving phase-separation ^13,14^. Recent studies have investigated in more detail the biophysical characteristics of crucial SG proteins such as Pab1 ^11^ and Pub1 ^12^. Those analyses have identified several protein regions or domains important for LLPS, and a more detailed definition of these protein hot spots is desired for a deeper understanding of the process. However, similar studies of other key SG components (e.g. eIF4G) are lacking.

SG are reservoirs for inactive mRNAs that are likely paused at their translation initiation stage ^5,15-18^. SG can exchange materials with P-bodies, for mRNA degradation and recycling, or can serve as platforms for restarting translation when the stress has ceased. In the latter scenario, it is possible that some of the multivalent interactions in the condensates are common to translation initiation complexes, providing structural links between SG nucleation and regular translation. Protein synthesis is tightly regulated by different mechanisms and signaling networks that involve dozens of proteins ^19-21^. Translation initiation is the rate-limiting step and is enhanced by mRNA 3’/5’-end circularization, which can be achieved by formation of a “closed-loop” complex (CLC) ^20^. In this model, Pab1 interacts with both the 3’ poly(A) tail and with eIF4G, a component of the eIF4F heterotrimer (eIF4G+eIF4E+eIF4A) that recognizes the 5’-cap (via eIF4E).

eIF4G is therefore a central player in both transcription initiation ^22,23^ and in SG nucleation ^3-5^ and is thus an attractive target for study of the link between these two processes. There are two eIF4G genes in *Saccharomyces cerevisiae* ^24^ and the most abundant one - eIF4G1 (also referred to as Tif4631) - contains several predicted IDRs, the longest one spanning residues 1-400. This region is important for eIF4G1 regulation and contains the binding sites for Pab1 ^25,26^, Pub1 ^27^ and RNA ^28,29^. The C-terminal part of eIF4G1 includes the domains for interaction with eIF4A ^30^ and eIF4E ^31^, two more RNA binding regions ^28,29^, and is also involved in interactions with translational repressors ^32^.

The highly abundant RNA-binding proteins Pab1 and Pub1 contain four and three RRMs (RNA Recognition Motifs) respectively that are involved in their interaction with the N-terminal eIF4G1 IDR ^22,25,27^, and they also contain various low complexity domains (LCD). These proteins are constituents of biological condensates and, in response to temperature increase or acidic pH ^10-12^, undergo *in vitro* LLPS in which both the RRMs and the LCD participate.

We here report NMR structural characterization of the eIF4G1 N-terminal IDR, as well as its interactions with Pab1 and Pub1, which reveals new insights into the roles of these three proteins in translation initiation and stress granule assembly. Our thorough NMR study provides a conformational ensemble of this eIF4G1 IDR that is stabilized by cation-π and π-π interactions. We also investigated self-recognition processes of Pab1, Pub1 and eIF4G1, and identified their mutual interaction sites. We propose models in which these multivalent interactions provide novel understanding of the different mechanisms of translation initiation and assembly of multiprotein condensates in SG.

## RESULTS

### The N-terminal eIF4G1 IDR contains residual structural features

Although the N-terminal region of *Saccharomyces cerevisiae* eIF4G1 is predicted to be intrinsically disordered, it contains three boxes of about 15-20 conserved residues each, and a conserved RNA binding region (RNA1), within the first 249 amino acids ^28^ (Figure 1A). We studied this eIF4G1 region using an eIF4G1_1-249_ construct that is stable for days under different pH conditions (Supplementary Figure 1). We then analyzed eIF4G1_1-249_ using NMR, which is a very powerful technique for investigation of IDRs and their interactions at the residue level (^33,34^ and references therein).

**Figure 1.**
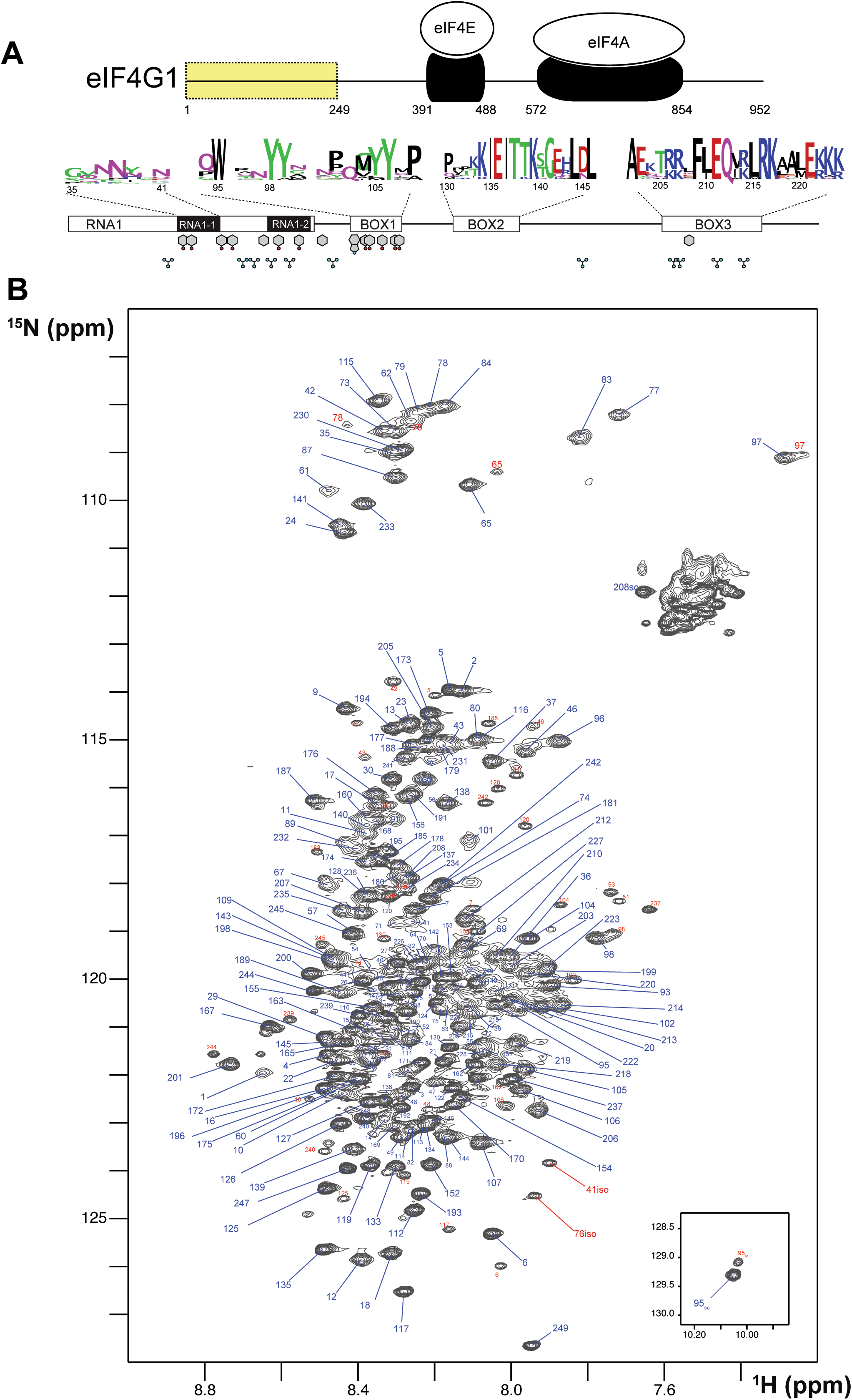
Sequence features and NMR spectrum of eIF4G1_1-249_. **(A)** Schematic representation of *S. cerevisiae* eIF4G1 showing the interaction domains for the other two components of the eIF4F heterotrimer (in black). The N-terminal region containing the conserved boxes (residues 1-249) is highlighted in yellow. The conserved features (BOXes) of eIF4G1_1-249_ are represented below including weblogos indicating the conservation of each box across *Saccharomyces* and the position of key side chains (Tyr, Phe, Trp and Arg) capable of π-π and π-cation interactions. **(B)** ^1^H-^15^N HQSC spectrum of eIF4G1_1-249._ Each assigned residue is labeled: blue, major form; red, minor species.

The ^1^H-^15^N HSQC NMR spectrum of eIF4G1_1-249_ is characteristic of an intrinsically disordered protein (IDP) with low dispersion in the proton dimension and sharp signals (Figure 1B). However, interestingly, several glycines showed relatively broad peaks at δ_NH_ < 8.00 ppms (e.g. G97, G77 and G83) that are not compatible with a fully disordered state. We identified several minor species (signals labelled in red) that correspond to *cis*Pro conformers, and two uncommon chemical isomerization forms at positions 41 and 76 that were assigned to isoaspartates (Supplementary Figure 2). These variants arise from deamidation of N41 and N76, that lie next to Gly residues in the protein sequence; Asn-Gly sequences are known to have the highest tendency to experience this non-enzymatic deamidation in model peptides ^35^. The level of deamidation is similar for the two positions (12-14%) and remained constant in different samples and over NMR experimental time, suggesting that these forms might have been generated *in vivo*.

Analysis of eIF4G1_1-249_ secondary structure based on ^13^C chemical shifts, T_1_/T_2_ ^15^N relaxation times and residual dipolar couplings (RDCs) revealed the presence of an α-helix within BOX3 (Figure 2A). This finding was confirmed by characteristic sequential amide-amide NOEs measured in a 3D ^1^H-^15^N-HSQC-NOESY-^1^H-^15^N HSQC spectrum (Supplementary Figure 3). We determined the NMR structure of this α-helix using a BOX3 model peptide (eIF4G1_187-234_) (Supplementary Figure 3). No further standard secondary structure elements were identified in eIF4G1_1-249_. However, the broad glycine peaks seen in Figure 1B, suggested the presence of residual higher order structures in this construct. In support of such structures, random coil index” (RCI) values S2 predicted from the chemical shifts ^36^, and the lower T_1_, T_2_ relaxation times for the conserved boxes suggested that these boxes might be involved in transient contacts that restrict mobility and/or induce chemical exchange processes resulting in short T_2_ values (Supplementary Figure 4). The existence of long-range interactions was evidenced by paramagnetic relaxation enhancement (PRE) measurements. The nitroxyl spin-label derivatization of engineered cysteine mutants (eIF4G1_1-249_ has no native Cys) indicates long-range PREs for S200C and Q109C mutants (Figure 2B). In a protein such as eIF4G1_1-249,_ the PREs (calculated as described in ^37^ and Supplementary Figure 5) are expected to occur within 25-30 Å of the spin label. Therefore, the PRE data suggested the presence of long-range contacts in eIF4G1_1-249_ involving BOX1, RNA1 and BOX3. To determine if these contacts are predominantly intra- or intermolecular, we placed the spin label in the non-isotope labeled Q109C mutant, added wild-type ^15^N-labeled eIF4G1_1-249_, and then measured PRE. The PRE fingerprint shows that eIF4G1_1-249_ can dimerize/oligomerize through contacts involving BOX1 and RNA1, as these elements “sense” the presence of the spin label in *trans* (Figure 2B lowest graph). However, the magnitude of the effects of the spin label in *trans* is smaller than when it is in *cis* (Figure 2B middle graph), suggesting that there is a small population of the self-associated species. All of the Tyr residues of the construct are contained in these regions involved in eIF4G1_1-249_ self-recognition (see Figure 1A). Five of these Tyr resides are included in BOX1 that is a predicted pro-amyloidogenic sequence (Supplementary Figure 6). These data suggest that dimerization/oligomerization of eIF4G1_1-249_ involves Tyr-Tyr interaction and perhaps leads to early amyloid-like stages.

**Figure 2.**
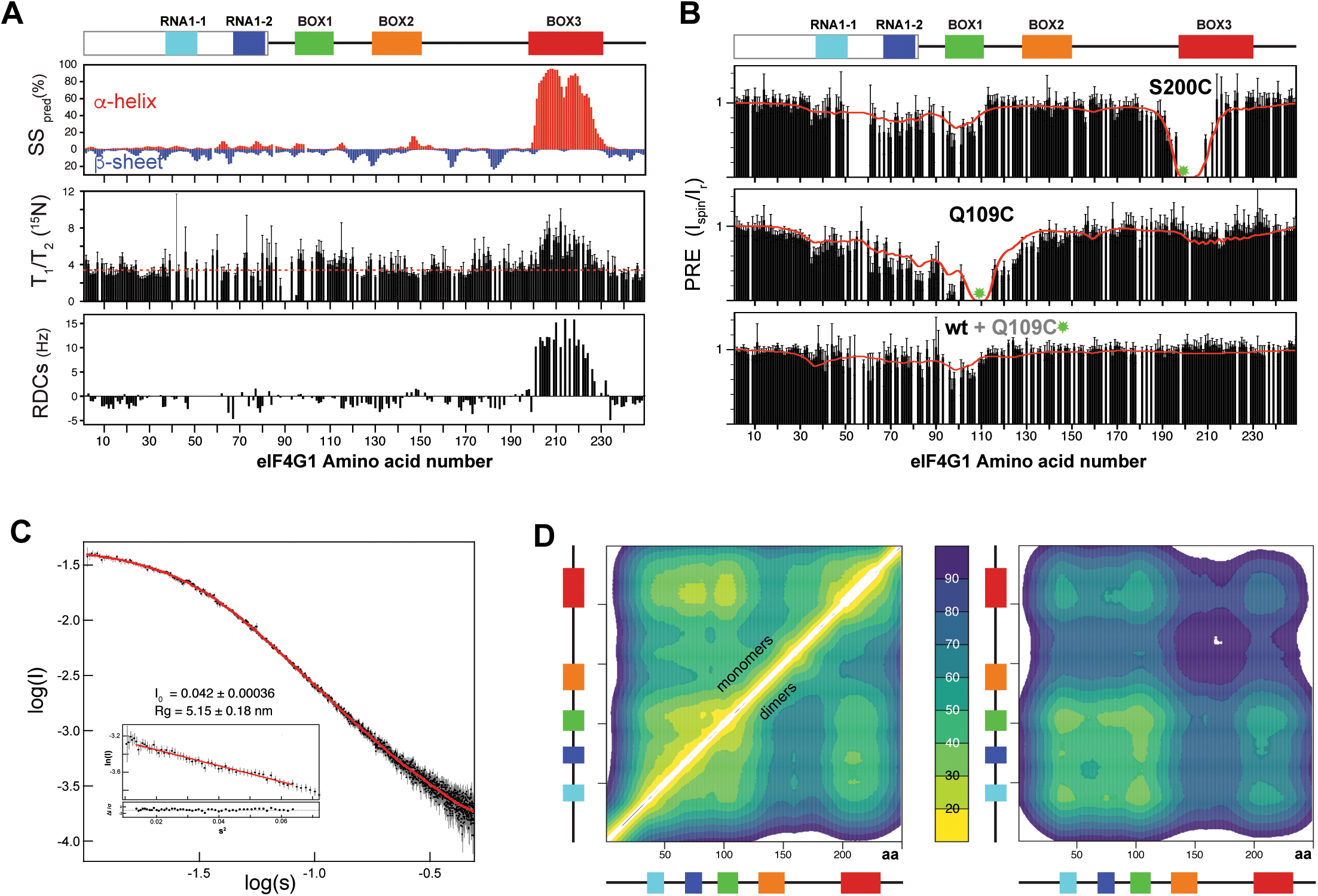
NMR structural analysis of eIF4G1_1-249_. **(A)** NMR evidence of residual secondary structure (per residue): percentage of predicted secondary structure (SS pred) calculated using the d2D program ^36^ (upper graph), ^15^N relaxation T_1_/T_2_ (middle graph, dotted red line marks the average) and residual dipolar couplings (RDCs) (lower graph). Conserved sequence elements in eIF4G1_1-249_ are represented at the top of the figure. (**B)** Per residue effect of paramagnetic relaxation enhancement (PRE) over relative signal intensities of two individual ^15^N-labelled eIF4G1_1-249_ mutants and of ^15^N-labelled wild-type eIF4G1_1-249_ mixed (1:1) with spin-labeled eIF4G1_1-249_ Q109C at natural isotopic abundance. Green circles mark the position of spin-labels. The red lines indicate the back-calculated PRE effects across the 500-member ensembles of monomers (upper and middle graphs) and dimers (lower graph). **(C)** Experimental SAXS curve showing log of intensity (I) versus log of scatter (S) of eIF4G1_1-249_ and EOM fitting obtained from the eIF4G1_1-249_ atomic models (red curve). Inset: Guinier analysis and the derived Radius of gyration (*R*_*g*_) and forward scattering intensity (*I(0)*) values. **(D)** (left panel) Ensemble-averaged intramolecular Cα contact maps obtained from the monomers and dimers (upper and lower triangles, respectively), and (right panel) intermolecular Cα contact maps. The average distances (in Å) were color-coded according to the scale in the middle, and the conserved boxes of eIF4G1_1-249_ are indicated along the axes of each map.

### eIF4G1 IDR conformational ensemble

IDPs, are considered to consist of ensembles of co-existing conformers. To build realistic conformational ensembles of IDPs, it is necessary to identify the possible residual secondary structures and long-range interactions between different regions of the polypeptide ^38^. These structural features are generally sparsely populated and transient in IDPs, which makes them difficult to detect and quantify experimentally thereby undermining the possibilities of calculating conformational ensembles. Nevertheless, several studies have proposed that interactions that favor aggregation, flexibility and/or long-range contacts are prevalent in IDPs ^8,39-41^. In particular, a recent report emphasized the key role of cation-π and π-π interactions in the “molecular grammar” of phase separation in prion-like IDPs ^39^. There are 11 Arg, 11 Tyr, 3 Phe and 1 Trp in eIF4G1_1-249_ that are suitable for these types of interactions, and they are mostly located in the conserved boxes (see Figure 1A). We therefore hypothesized that cation-π and π-π interactions (involving Arg, Tyr, Phe and Trp) might dominate the long-range contacts in eIF4G1_1-249_.

Different methods have been used to calculate IDP conformational ensembles using experimental data (NMR, SAXS and others) and/or computational approaches ^42-47^. Here we used the algorithm in the program Cyana 3.0 ^48^ for fast generation of eIF4G1_1-249_ all-atoms structural models. This approach allows a straightforward implementation of: (1) NOE-derived distance restraints and ^13^C-derived φ/ψ dihedral restraints for the parts of the protein that are well-folded (i.e. BOX3) and (2) ambiguous restraints for the cation-π and π-π interactions (involving Arg and Tyr) that we propose as dominant transient residue-residue contacts. We refer to these latter types of restraints as “knowledge-based” and used cautions to avoid biases in their selection (see materials and methods for specific details). We calculated 80000 eIF4G1_1-249_ structures and sorted them using our own greedy algorithm that optimized the fitting to experimental PRE data stepwise. As a seed for the ensemble the protocol chooses the structure with minimum PRE violations and then continues building up the ensemble stepwise using the same criteria (i.e. incorporating the structure that, together with the previously selected structures(s), minimizes violations). The target function reached a minimum value relatively quickly and increased slowly afterwards (Supplementary Figure 7). We arbitrarily selected a final 500-member ensemble to ensure sufficient structural variability while still maintaining good agreement between the back-calculated and the experimental PREs (red line in Figure 2B). Because the intermolecular PRE data showed self-association, we performed a similar protocol for analysis of eIF4G1_1-249_ dimers, hypothesizing that Tyr-Tyr interactions are the driving force of dimerization. The SAXS curve evidenced the IDP nature of eIF4G1_1-249_ (Figure 2C). To validate the eIF4G1_1-249_ ensemble models, we applied the Ensemble Optimization Method (EOM) genetic algorithm ^49,50^ to model the SAXS curve. Pools for monomeric and dimeric conformations were used without restricting the relative percentages of each set. We performed 10 independent EOM calculations, and each of them resulted in excellent fittings of the experimental curve (Figure 2C). Importantly, similar calculations done with either eIF4G1_1-249_ monomers or dimers alone resulted in worse fits. In the SAXS-selected eIF4G1_1-249_ ensemble, the monomers dominate (88%). As expected, the back-calculated PREs from the collection of EOM ensembles showed poorer agreement with the experimental data, but neatly reflected the overall trends regarding the long-range contacts in eIF4G1_1-249_ (Supplementary Figure 7).

The eIF4G1_1-249_ ensemble of conformers showed a flexible α-helix in BOX3 and, despite the presence of long-range contacts, no predominant tertiary fold was found. The average Cα-Cα distance maps revealed that local and long-range contacts were prevalent between BOX1 and RNA1-1/RNA1-2 boxes, and between these three elements and BOX3 (Figure 2D left). In contrast, there was a remarkable absence of long-range interactions involving BOX2. Dimerization contacts were dominated by BOX1 and to a lesser extent by the RNA1-1 box (Figure 2D right). The BOX3 region also showed a minimum in the intermolecular Cα-Cα distance maps, due to the coexistence of intramolecular Arg-Tyr and intermolecular Tyr-Tyr contacts (Figure 2D right). Indeed, these π-π and cation-π interactions tend to appear in networks, rather than in binary mode, probably favored by the planar nature of aromatic and guanidinium groups.

In summary, these data showed that eIF4G1_1-249_ is predominantly disordered except for an α-helix in BOX3. Atomistic models were constructed with experimental and knowledge-based restraints and ensembles were built by restraining against experimental NMR and SAXS data. Their analysis showed an intrinsic tendency of eIF4G1_1-249_ to dimerize (oligomerize), in which BOX1 plays the chief role.

### Pub1 and Pab1 self-recognition

eIF4G1 interacts with RBPs including Pub1 and Pab1 that are key components of SG ^5,6^. These two proteins have a similar architecture, with 3 and 4 RRM domains, respectively, and various IDRs (Figure 3A&B), and both are involved in LLPS ^11,12^. We studied Pub1 and Pab1 constructs using NMR and SAXS (Figure 3) to characterize homotypic interactions and self-assembly processes.

**Figure 3.**
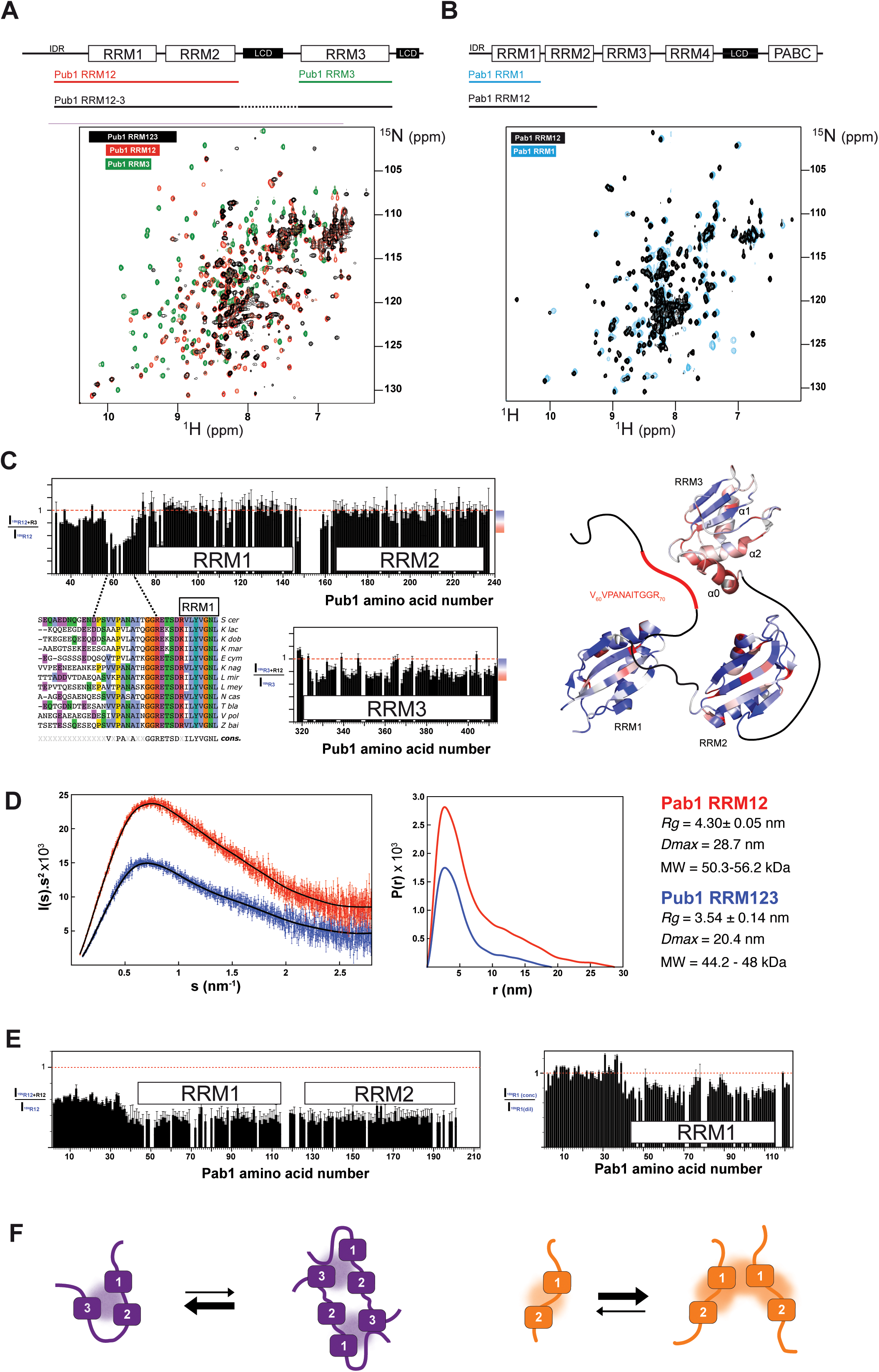
NMR study of Pub1 and Pab1 self-recognition. Multidomain architecture and superposition of the NMR ^1^H-^15^N HQSC spectra of the indicated Pub1 (**A**) and Pab1 (**B**) constructs. Signals were color-coded according to the specific construct. (**C**) Intermolecular interactions between Pub1 RRM12 and Pub1 RRM3 monitored as the change in HSQC signal intensities upon titration of one ^15^N-labeled construct with another unlabeled construct. Intensity variations are mapped onto the structural model of Pub1 RRM123 according the color-scale at right of the graphs. The folded domains correspond to the structures of Pub1 RRM12 (PDB:3MD3) and Pub1 RRM3 (PDB:2LA4) and the interconnected segments (IDRs) are indicated as lines. Red areas represent hot spots involved in transient contacts. The sequence alignment of different yeast Pub1 homologs is shown for the segment that precedes RRM1. Dashed lines show the limits of the region involved in transient contacts with RRM3. Conserved residues are colored by amino acid type and the consensus sequence (***cons***.) is shown below the alignment. (**D**) Kratky plots (right hand panel) of the SAXS experimental curves for Pab1 RRM12 (red) and Pub1 (RRM123) (blue). The continuous lines in the right hand panel show the fitting corresponding to the distance probability distributions (*P(r)*). Relevant hydrodynamic parameters obtained by Guinier analysis: Radius of gyration (*Rg*) and maximum particle dimension (*D*_*max*_) are shown in the table at right together with the predicted molecular weight ranges using the Bayesian approach ^51^. (**D**) (left) Pab1 RRM12 self-association monitored by changes in the HSQC signal intensities ratios between an ^15^N-Pab1 RRM12 (200 µM) sample and an ^15^N-Pab1 RRM12 (200 µM) + Unlabeled Pab1 RRM12 (200 µM) (1:1) sample. Thus the concentration of ^15^N-Pab1 RRM12 (NMR visible) was 200 µM in each of the samples, whereas the total Pab1 RRM12 concentration was increased to 400 µM in the second sample. The two RRMs display higher reduction in intensities in comparison with the N-terminal IDR (aa 1-35). The right panel shows the ratios of signal intensities of ^15^N HSQCs measured at two different concentrations (35 µM and 200 µM) of ^15^N-Pab1 RRM1. Absolute intensities of the dilute sample were scaled up prior to comparison. (**F**) Proposed structural models of interdomain contacts in monomeric and dimeric forms of Pub1 RRM123 (purple) and Pab1 RRM12 (orange) based on NMR and SAXS analyses. Diffuse clouds represent transient interactions.

^15^N relaxation data for Pub1 RRM12 and Pub1 RRM3 constructs showed that, as expected, the RRM domains have slower mobility than the N-terminal IDR of Pub1 RRM12 (Supplementary Figure 8A). The correlation times (τ _c_) derived from these data showed that Pub1 RRM3 behaves as an 11 kDa globular protein, whereas RRM12 exhibits slower tumbling, although it is not as slow as that of a rigid RRM1-RRM2 tandem (∼20 kDa) (Supplementary Figure 8B). This analysis suggested that the RRMs in the Pub1 RRM12 and Pub1 RRM3 constructs essentially behave as dynamically independent modules (somewhat constrained by their connection through the interdomain linker for Pub1 RRM12). In contrast, the above-described Pub1 RRM123 ^1^H-^15^N HSQC spectrum (see Figure 3A, in black) showed substantial line broadening compared with that of RRM12 or RRM3, suggesting that the mobility of the RRMs is more restricted in this construct, likely due to interactions between the RRMs. To investigate such interactions, we performed NMR titration analysis of Pub1 RRM12 and RRM3. Chemical shift perturbations of the Pub1 RRM3 spectrum were small, but the ratios of the signal intensities of ^15^N Pub1 RRM3 and ^15^N Pub1 RRM3+Pub1 RRM12 samples evidenced transient interactions that can be mapped to helices 0, 1 and 2 (Figure 3C lower graph and mapping on the structure). The reverse titration highlighted the Pub1 RRM12 residues involved in transient contacts, which are located in a short and conserved hydrophobic segment (V_60_VPANAITGGR_70_) within the Pub1 N-terminal IDR and in discrete spots in the RRMs (Figure 3C upper graph, data mapping on the structure and sequence alignment).

**Figure 4.**
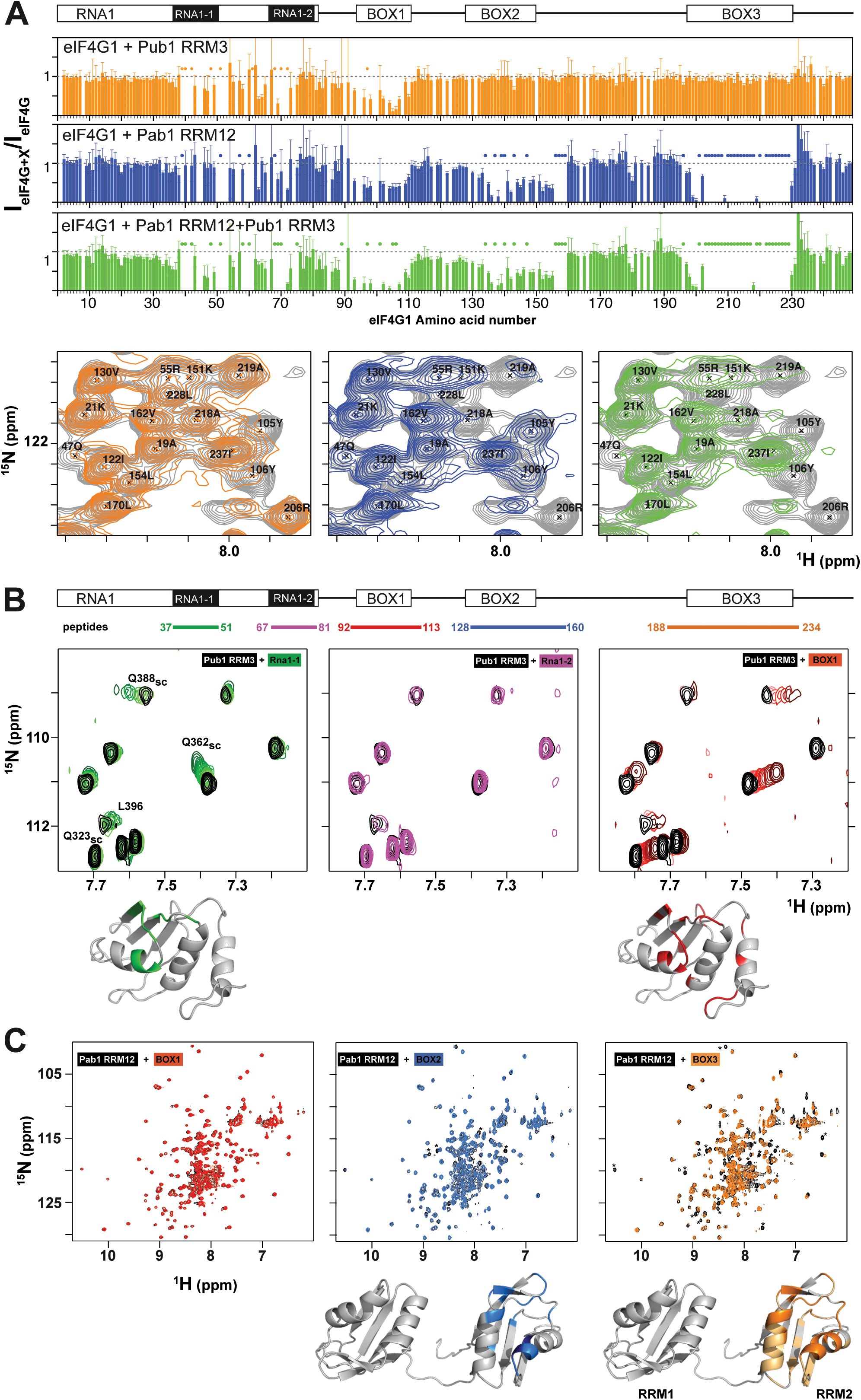
Mapping of Pub1 and Pab1 binding sites on eIF4G1_1-249_. **(A)** Sequence dependence of eIF4G1_1-249_ ^1^H-^15^N HSQC signal intensity ratios between free eIF4G1 _1-249_ and various eIF4G1_1-249_ complexes with Pub1 RRM3 (orange), Pab1 RRM12 (blue) and Pub1 RRM3+Pab1 RRM12 (green). A representative region of the eIF4G1_1-249_ ^1^H-^15^N HSQC spectrum (in grey) is shown below superimposed with the equivalent spectra of the Pub1 RRM3 (in orange), Pab1 RRM12 (blue) and Pub1 RRM3+Pab1 RRM12 (green) complexes. Specific residues were labelled to illustrate the selective disappearance of eIF4G1_1-249_ signals upon complex formation in each case (marked with dots in the bar charts). **(B, C)** NMR study of the interaction of eIF4G1 peptides with Pub1 RRM3 **(B)** and Pab1 RRM12 **(C)** monitored on their ^1^H-^15^N HSQC spectra. The regions of eIF4G1_1-249_ that correspond to each of the five peptides tested are indicated at the top in (B). Black: spectra of the free proteins; Colors: peptide titrations according to the color scheme in (B). Chemical shift perturbations and signal broadening were mapped onto the model structures of Pub1 RRM3 (PDB:2LA4) and Pab1 RRM12 (bottom).

**Figure 5.**
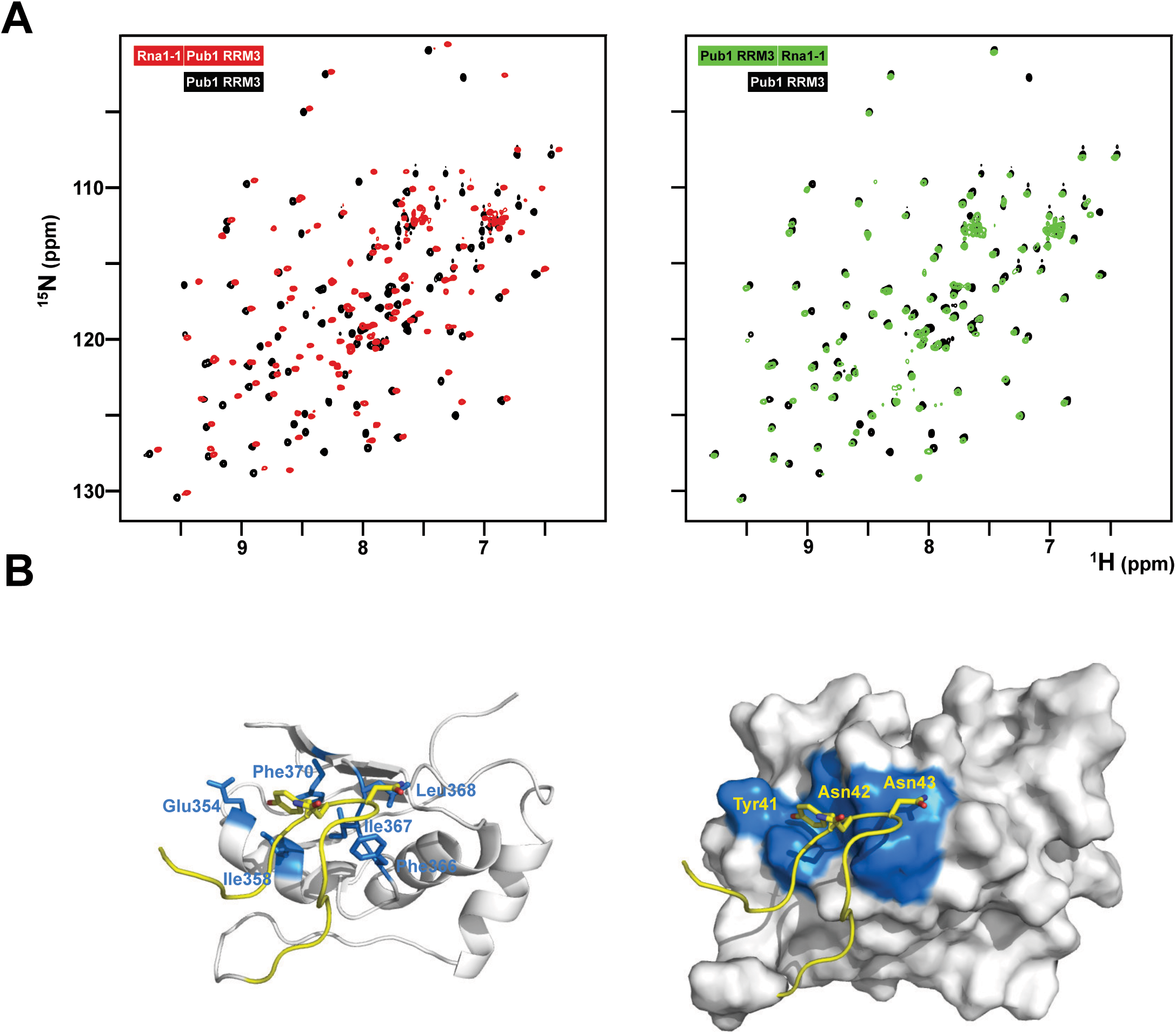
Structural characterization of the eIF4G1-Pub1 interaction. **(A)** Superposition of 2D ^1^H-^15^N HSQC of free Pub1 RRM3 (black) on either eIF4G1_35-49_-Pub1 RRM3 (red) or Pub1 RRM3-eIF4G1_35-49_ (green) chimeric constructs. **(B)** NMR structure of eIF4G1_35-49_-Pub1 RRM3. Yellow, eIF4G1; White, Pub1 RRM3; Blue, Pub1 RRM3 residues contacting eIF4G1. Key interacting residues are labelled on both eIF4G1 and Pub1RRM3 regions of the chimera.

**Figure 6.**
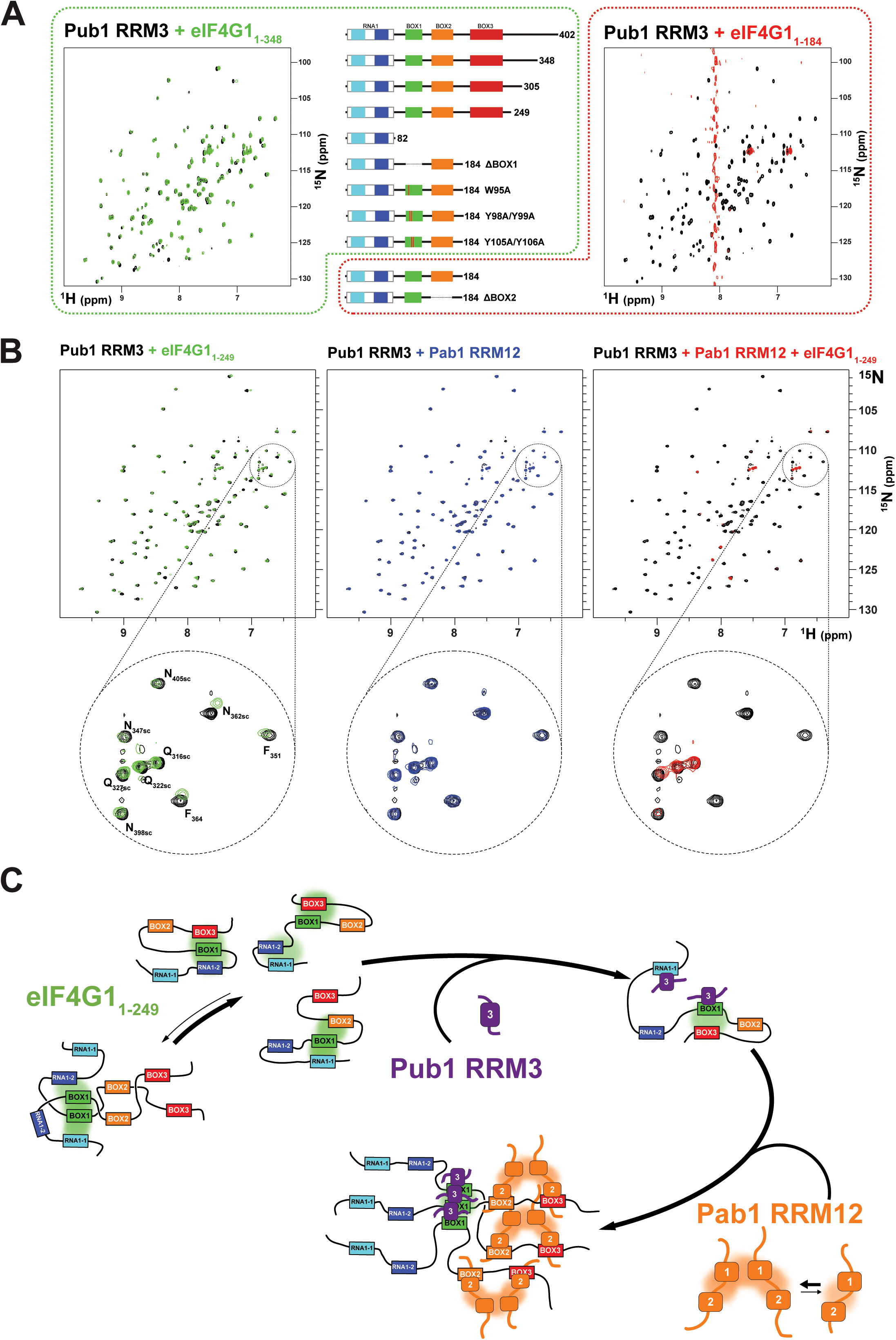
Pub1 RRM3 can interact with eIF4G1 through two different modes. **(A)** Superposition of the ^1^H-^15^N HSQC spectra of free Pub1 RRM3 (black) and Pub1 RRM3 in complex with eIF4G1_1-348_ (green) or eIF4G1_1-184_ (red). The Pub1 RRM3 NMR crosspeaks show small perturbations in the first complex (left panel) and completely disappear (with the exception of the highly mobile Asn/Gln side chain peaks) in the spectrum of the complex (right panel). The central panel shows eIF4G1 mutants that had similar effects as eIF4G1_1-348_ (outlined in green dash) or eIF4G1_1-184_ (outlined in red dash) on the Pub1 RRM3 spectrum. **(B)** Study of the effect of eIF4G1_1-249_ and/or Pab1 RRM12 titrations on the Pub1 RRM3 ^1^H-^15^N HSQC spectrum. Superposition of the Pub1 RRM3 spectra before (black signals) and after titration with eIF4G1_1-249_ (left, green signals), Pab1 RRM12 (middle, blue signals) and eIF4G1_1-249_ + Pab1 RRM12 (right, red signals). A small area of each spectrum is expanded below for a more detailed view. The Pub1 RRM3 signals are unperturbed upon titration with Pab1 RRM12 (blue spectrum), but disappear when Pab1 RRM12 is combined with eIF4G1_1-249_ (red spectrum). (**C**) A dual key interaction model between eIF4G1_1-249_, Pub1 RRM3 and Pab1 RRM12 to explain oligomerization. Multiple intramolecular interactions between BOX1 and other conserved elements of eIF4G1_1-249_ maintain it predominantly monomeric. Weak interactions with Pub1 RRM3 (first key) cancel out some of these transient contacts, but some remain (e.g. BOX1-BOX3) preventing BOX1-driven oligomerization. The interaction with Pab1 RRM12 (second key) further blocks internal contacts to BOX1 triggering its aggregation. Pub1 RRM3 binding to BOX1 oligomeric forms would be reinforced by Pub1-Pub1 interactions.

**Figure 7.**
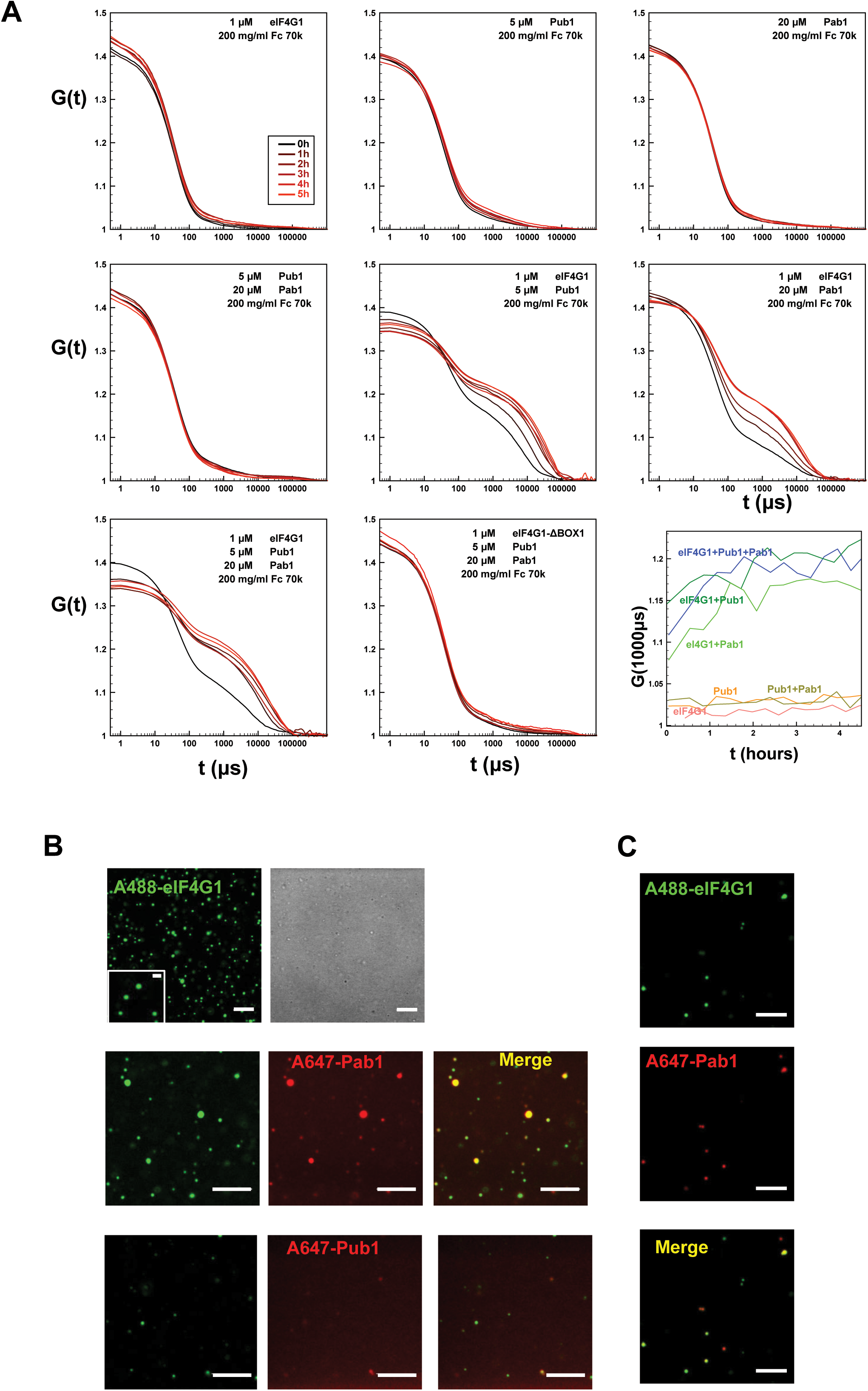
Formation of condensates by Pub1/Pab1/eIF4G1 mixtures in crowding conditions. **(A)** DLS analysis of the indicated individual Pub1 (Pub1 RRM123), Pab1 (Pab1 RRM12), and eIF4G1 (eIF4G1_1-249_ or eIF4G1_1-249_ ΔBOX1) proteins, and of their double and triple mixtures at the concentrations indicated in the figure in the presence of Ficoll (Fc) 70k. Time-dependence (in µs) of the autocorrelation functions G(t) is shown for different protein mixtures and recorded a different time after mixing (panel code shown in the upper left panel). The lower right panel (row 3, column 3) shows the time evolution of the autocorrelation function at 1000 µs for different mixtures of previous graphs, that correspond to the second phase associated to the aggregates. **(B)** Representative confocal fluorescence microscopic images of ternary mixtures eIF4G1_1-249_ (eIF4G1) labelled with Alexa 488 dye (A488) and Pab1 RRM12 (Pab1) plus Pub1 RRM123 (Pub1). In this ternary mixture, Pab1 plus Pub1 are either unlabeled (row 1, second column) or one of them is labeled with Alexa 647 dye (A647) as indicated in the figure (2^nd^ and 3^rd^ rows), while the other is unlabeled. **(C)** Confocal images of the mixture of eIF4G1_1-249_ (eIF4G1) and Pab1 RRM12 (Pab1), labelled with Alexa 488 and Alexa 647, respectively. In B and C, when present, the final concentration of eIF4G1_1-249_, Pab1 RRM12 and Pub1 RRM123 was 1, 20 and 5 μM, respectively. The concentration of labelled proteins was 1 μM and additional unlabeled protein was added to achieve the indicated final concentration. Scale bars, 5 μm. Inset scale bar, 1 μm. In A, B and C, samples contained Ficoll 70 (200 g/L) as a crowding agent.

**Figure 8.**
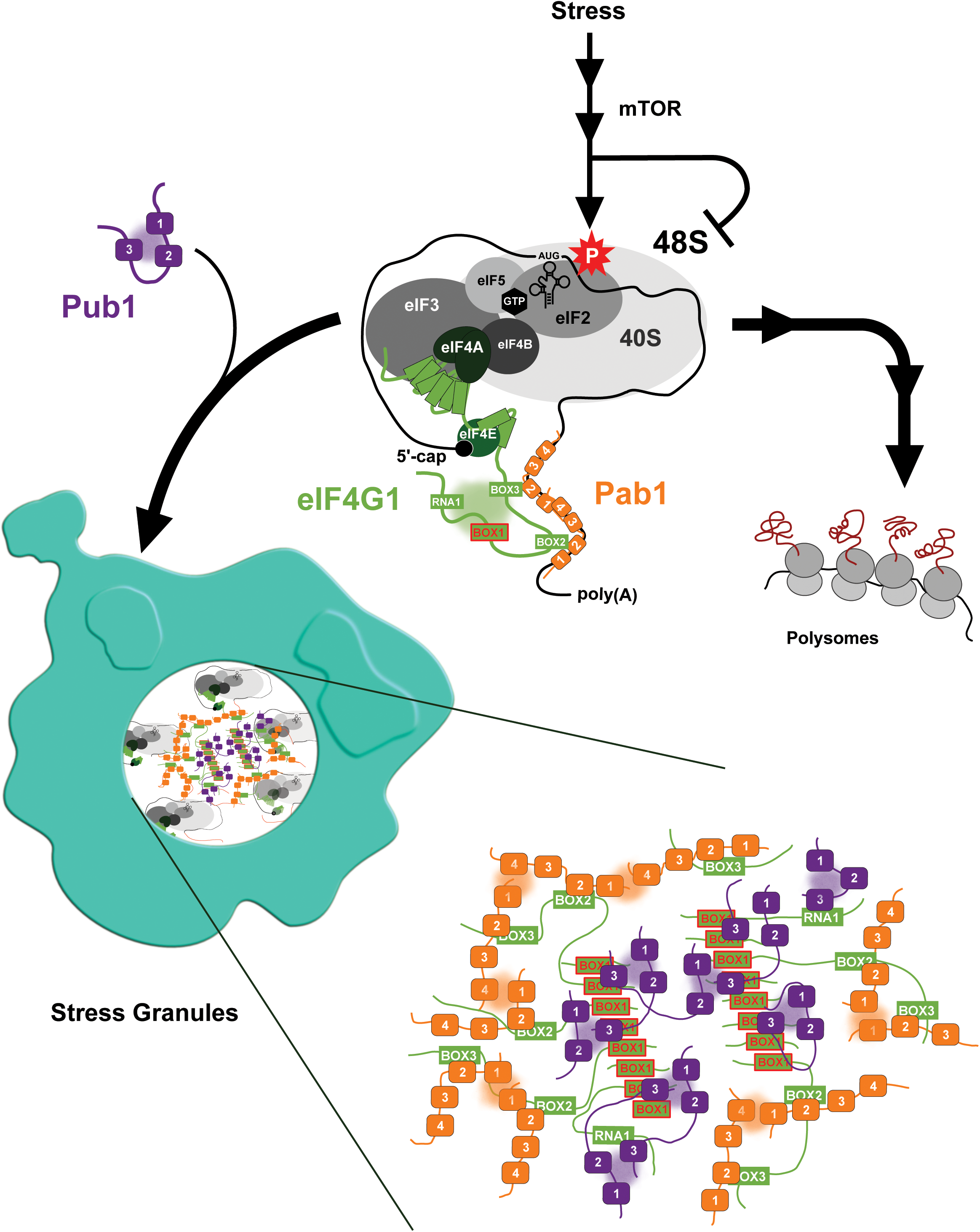
Simplified model of the stress response highlighting the role Pab1/Pub1/eIF4G1 multivalent interactions. Phosphorylation of eIF2alpha by the integrated response signaling pathway block translation in the 48S pre-initiation complex. Components of the complex have been represented schematically in different shades of grey and eIF2alpha phosphorylation is represented by a red star. The mRNA is represented by black line with a circle representing the 5-end capping, recognized by eIF4E, and the 3’-end poly(A) tail bound by Pab1 (in orange) (for simplicity only RRMs have been represented). eIF4G1 has been colored in green and transient interactions between the different boxes in the N-terminal IDR, that help to stabilize it in a monomeric state, are shown as a green cloud. Pub1 (in purple) would interact with RNA1_1 and BOX1 elements triggering eIF4G1 aggregation through BOX1 and later the assembly of the SG. The composition of the SG has been greatly simplified to show only the multivalent interactions between Pub1, Pab1, eIF4G1. The expanded view shows details of an structural model of the SG cores that incorporate these interactions. The model excludes other homotypic (involving other SG proteins) and heterotypic (with RNA) interactions for simplicity.

We investigated these interactions in the Pub1 RRM123 construct using SAXS. The data show the characteristic profile of a multidomain protein, and the Radius of gyration (*R*_*g*_), the pair-wise distance distribution, *P(r)*, and Bayesian inference molecular weight analysis ^51^ suggested that the protein is mainly monomeric (Figure 3D) at the studied concentration, although a small contribution of dimers cannot be discounted (Pub1 RRM123 theoretical MW is 35 kDa). In agreement with this conclusion, fitting of the SAXS curve using EOM was better with monomers, rendering a variety of reasonable compact models with interactions among the RRMs (Supplementary Figure 9). In general, both NMR and SAXS data support a model of transient contacts between the RRMs (e.g. RRM3 to RRM1/2) and between the RRMs and the IDR regions of Pub1 RRM123. Such contacts could, under certain circumstances, occur intermolecularly, thereby promoting oligomerization (Figure 3F left).

Interestingly, a similar arrangement involving intramolecular cyclic species and dimers has been proposed for yeast Pab1 ^52^, and recently intermolecular contacts between Pab1 RRM1 and RRM4 have been shown in the context of poly(A) RNP ^53^. Here, we assigned the Pab1 RRM12 ^1^H-^15^N HQSC spectrum (Figure 3B in black and Supplementary Figure 10). Excluding the N-terminal IDR (residues 1-35), Pab1 RRM12 showed large T_1_/T_2_ ratios. The calculated correlation times (τ_c_) were higher than expected for a single rigid RRM-RRM tandem but not as high as that for a globular dimer (Supplementary Figure 8B). This dynamic behavior was in contrast with that of Pub1 RRM12, a molecule of similar size and architecture (Supplementary Figure 8). Titration of ^15^N Pab1 RRM12 with unlabeled Pab1 RRM12 showed that there was a selective reduction in signal intensities of the RRM12 tandem (Figure 3E left). Similar reduction in the RRM was observed when comparing normalized intensities at two concentrations of the Pab1 RRM1 construct (Figure 3E right). On the other hand, the SAXS data strongly suggested that Pab1 RRM12 is predominantly dimeric (Figure 3D), with a molecular weight range close to the theoretical value of the Pab1 RRM12 dimer (49 kDa). Although the EOM analysis of the SAXS data required dimerization it cannot discriminate between whether it is driven by RRM1 or RRM2 (Supplementary Figure 9). However, given that the spectra of Pab1 RRM1 and Pab1 RRM12 were highly superimposable (see Figure 3B), we propose that RRM1 is the dimerization domain, as previously suggested ^52 54,55^ (Figure 3F right). In this model, transient intramolecular contacts between RRM1 and RRM2, perhaps similar to those reported between RRM1 and RRM2 in complex with RNA ^53-55^, would explain why RRM2 exhibited similar dynamics (Supplementary Figure 8) and concentration dependent responses (Figure 3E left) as RRM1.

To summarize these results, both Pub1 and Pab1 can form dimers to a different extent, involving specific interactions between RRMs and IDRs that are expected to be important in quinary structures ^56^ that include these molecules.

### eIF4G1 interacts with Pab1 and Pub1 through multiple binding sites

After characterizing Pab1 and Pub1 individually, we next studied the structural details of their interactions with eIF4G1_1-249_. Previous to this work, we showed that Pub1 RRM3 interacts with eIF4G1 ^27^. Here we mapped this interaction on eIF4G1_1-249_ by analysis of changes on the ^1^H-^15^N HSQC spectra, which indicated three putative binding sites for RRM3 in eIF4G1_1-249_ (orange bar chart in Figure 4A): two in RNA1 (RNA1-1 and RNA1-2) and one in BOX1. Strikingly, these sites have a small consensus sequence motif (YNNxxxY), only present in this region of the eIF4G1. We tested the ability of short peptides of eIF4G1 that corresponded to the conserved elements (Figure 4B) to bind to ^15^N-labelled Pub1 RRM3, by monitoring their effect on the Pub1 RRM3 ^1^H-^15^N HSQC spectrum. Only BOX1 and RNA1-1 peptides caused changes in the Pub1 spectrum arising from direct contacts (Figure 4B). The Pub1 binding site in BOX1 overlaps with the pro-amyloidogenic sequence (Supplementary Figure 6). The absence of spectral changes upon RNA1-2 peptide titration (Figure 4B) suggested that the effects observed on the spectrum of the corresponding region of eIF4G1_1-249_ in complex with Pub1 RRM3 (orange bar chart in Figure 4A) are probably due to conformational rearrangements.

The binding of short eIF4G1 peptides was too weak to obtain structural restraints (e.g. intermolecular NOEs) for calculation of the structure of the complex. To overcome this technical problem, we constructed recombinant chimeras of eIF4G1_35-49_ and Pub1 RRM3 (Figure 5 and Supplementary Figure 11). The NMR spectrum of the eIF4G1 peptide fused to the C-terminus of Pub1 was similar to that of Pub1 RRM3 alone (Figure 5A right), whereas that of the N-terminally fused chimera differed significantly (Figure 5A, left), suggesting that the peptide can effectively fold-back into the binding site only in the latter case. Using this latter construct, we obtained enough experimental restraints to calculate the structure of the eIF4G1_35-49_-Pub1 RRM3 chimera, which shed light on the key elements required for molecular recognition (Figure 5B and Supplementary Figure 11). The structure showed that eIF4G1 residues Y_41_, N_42_ and N_43_ (part of the YNNxxxY conserved motif) interact with a shallow cleft in Pub1 RRM3, defined by the contact between helix α1 and strand β2. The eIF4G1 Y_41_ is inserted into a small cavity and contacts I_358_, I_367_ and F_370_ of Pub1, whereas the N_42_ and N_43_ of eIF4G1 are more exposed but contact L_368_ and F_366_ of Pub1. The latter residue was previously shown to be important for eIF4G1-Pub1 interaction ^27^. The interaction surface was small, in agreement with a weak eIF4G1-Pub1 interaction.

We next studied the interaction between eIF4G1 and Pab1 using similar approaches. NMR titrations of unlabeled Pab1 RRM12 over ^15^N-eIF4G1_1-249_ (blue bar chart in Figure 4A) caused perturbations and signal disappearance in three well-delimited regions: aa 95-103 (BOX1), aa 135-160 (BOX2) and aa 200-234 (BOX3). These are very important results since, to date, only BOX3 has been reported as a Pab1 binding site ^26^. As done for Pub1 RRM3, we studied the interactions of Pab1 RRM12 with elF4G1 fragments to validate the putative binding sites. As expected, the BOX3 peptide interacted with Pab1 RRM12 causing significant perturbations in the helix1-helix2 interface of RRM2 (Figure 4C right), a region equivalent to that involved in human eIF4G-PABP1 recognition ^55^. The BOX2 peptide also interacted with RRM2 through a similar interface but it probably only weakly interacts, because it caused fewer changes than the BOX3 peptide. NMR data showed that the BOX1 peptide does not directly interact with Pab1 RRM12 and that the observed changes in that region of eIF4G1_1-249_ (blue bar chart in Figure 4A) are due to reorganization of internal contacts. Interestingly Pub1 and Pab1 used different interfaces of the RRM to interact with eIF4G1 (Supplementary Figure 11C).

These results showed that, although Pub1 and Pab1 interact with eIF4G1_1-249_ through multiple sites, most of these interactions are weak because they caused minor chemical shift changes in these RBPs; the exception is the Pab1-BOX3 interaction that showed changes of a larger magnitude. The presence of two eIF4G1 binding sites for Pub1 and Pab1, combined with their self-association properties, might suggest a possible cooperative recognition mode of eIF4G1.

### eIF4G1 BOX1 controls Pub1 RRM3 oligomerization

During the course of the study, we found that Pub1 RRM3 interacts differently with different eIF4G1 constructs. Titration with eIF4G1_1-82_, eIF4G1_1-305_, eIF4G1_1-348_ and eIF4G1_1-402_ constructs caused small changes in the Pub1 RRM3 NMR signals equivalent to those described for its interaction with eIF4G1_1-249_ and BOX1 and RNA1-1 peptides (Figure 6A left). However, surprisingly, titration with eIF4G1_1-184_ cause the disappearance of nearly all of the Pub1 RRM3 ^1^H-^15^N HSQC crosspeaks (Figure 6A right). This result would be consistent with Pub1 RRM3 being part of a high molecular weight structure; probably eIF4G1-Pub1 oligomers. Nevertheless, we found that the origin of this differential binding mode resided in the BOX1 element of eIF4G1. Thus, Pub1 RRM3 bound weakly to several eIF4G1_1-184_ BOX1 mutants (ΔBOX1, W95A, Y98A/Y99A and Y105A/Y106A), whereas deletion of BOX2 (ΔBOX2) still caused the Pub1 aggregation-like pattern. Our hypothesis, based on these data is that Pub1 RRM3 might bind more efficiently to BOX1 oligomers, which are kept at a low population in the eIF4G1 free state due to interactions between BOX1 and other elements (Figure 2D). Interaction of Pub1 RRM3 with RNA1/BOX1 would switch off some of these intramolecular eIF4G1 interactions leading to eIF4G1_1-184_ oligomerization. However, Pub1 RRM3 binding would not disrupt the BOX3-BOX1 contacts present in longer eIF4G1 forms (eIF4G1_1-249,_ eIF4G1_1-305_, eIF4G1_1-348_ and eIF4G1_1-402_) thereby preventing the formation of high order species. To further test this theory, we investigated the effect of incorporating Pab1 RRM12 into the eIF4G1_1-249_/Pub1 RRM3 complex. In this ternary complex, the Pub1 RRM3 signals disappeared, suggesting that the Pab1 RRM12 binding to BOX2 and BOX3 acts as a second switch that releases the BOX1 oligomers. NMR data showed that Pub1 RRM3 did not interact with Pab1 RRM12 (Figure 6B middle panel). These data agree with the data described above in Figure 4A, where the presence of the Pab1+Pub1 mixture (Figure 4A, in green), caused larger BOX1 line broadening in the eIF4G1_1-249_ spectrum than the presence of either Pub1 or Pab1 alone (Figure 4A, in orange and blue, respectively), which also suggested the existence of BOX1-driven oligomers.

These NMR analyses suggested a two-key mechanism whereby Pub1 and Pab1 bind to eIF4G1 causing conformational changes that promote BOX1 self-assembly (Figure 6C). These two RBPs interact with eIF4G1 elements that contact with BOX1 in the free state (Figure 2D). These contacts likely prevent BOX1 aggregation, whereas the coordinated effect of Pub1/Pab1 binding enhances it.

### Pab1-Pub1-eIF4G1 form micrometer-size condensates

Previous studies showed that isolated Pub1, Pab1 and eIF4G1 proteins undergo LLPS upon pH and/or temperature stresses ^10-12^. LLPS is thought to be promoted by the RRMs of Pab1 ^11^ and Pub1 ^12^, although it has been suggested that the disordered regions could also be involved as regulatory elements. Those previous studies only analyzed individual proteins and did not explore the potential of homotypic interactions with other SG proteins to influence LLPS. Our NMR analysis suggested that simultaneous binding of Pab1 RRM12, eIF4G1_1-249_ and Pub1 RRM3 has the potential to form high order structures that cannot be detected by this technique because of their large size. To further investigate this possibility, we determined if different Pub1/Pab1/eIF4G1 combinations could form microscopic condensates that might resemble biological condensates. For such experiments we used protein concentrations similar to those used in the above-mentioned previous studies ^10-12^ and used Ficoll 70 (200 g/l) to simulate crowding in the cellular environment. Dynamic light scattering (DLS) indicated that large aggregates were formed in some protein combinations (Figure 7A). In contrast, the individual proteins exhibited autocorrelation functions that did not differ from that of Ficoll-70 alone, suggesting that the individual proteins did not aggregate. Moreover, the curve profiles of the single proteins remained stable for several hours. The Pab1:Pub1 mixture (Figure 7, row 2, column 1) showed the same behavior, but other double protein mixtures and the triple one showed a second phase, evidencing the presence of micrometer-size particles. These particles were present right from the beginning and appeared to reach a maximum within 2 hours of mixing (Figure 7A, row 3, column 3). Because, all of these combinations contained eIF4G1 and at least one RNA binding protein, we concluded that the interactions between eIF4G1 and Pab1/Pub1 promoted aggregation; probably by enhancing the intrinsic propensity of BOX1. To test this hypothesis we replaced eIF4G1_1-249_ by the eIF4G1 ΔBOX1 mutant and the triple mixture showed no aggregation (Figure 7, row 3, column 2). This result strongly suggested that the BOX1 element, a Tyr-rich pro-amyloidogenic region (Supplementary Figure 6), is necessary for the formation of high molecular weight aggregates. In support of this result, we found that eIF4G1 constructs containing BOX1, but not BOX1-delection mutants, could form hydrogels (Supplementary Figure 6).

Further insight into the DLS-detected particles was obtained using confocal fluorescence microscopy. Images of triple mixtures containing Pab1 RRM12, Pub1 RRM123 and Alexa 488 labelled eIF4G1_1-249_ in Ficoll 70 (200 g/L) revealed the presence of discrete rounded particles (∼1 µm and smaller, Figure 7 B upper panels). Both, Pab1 and Pub1 were observed to colocalize with eIF4G1 in these assemblies, as observed in fluorescent images in which the proteins were pairwise labelled with spectrally different dyes (Alexa 488 and Alexa 647, Figure 7B middle and lower panels). Similar structures were observed for binary Pab1/eIF4G1 mixtures (Figure 7C), in good agreement with the DLS measurements.

These results showed that the eIF4G1/Pab1/Pub1 mixtures could form crowding-driven structures resembling those previously described for full length Pab1 or Pub1 ^11,12^, but without the requirement for pH or temperature stress. The agreement of the DLS and fluorescence microscopy results with the NMR data suggested that Pub1, Pab1 and eIF4G1 can form heterologous oligomers of discrete size, that might represent early stages of protein condensates.

## Discussion

This study provides an extensive NMR characterization of eIF4G1_1-249_ at the residue level, its complex conformational landscape and its interactions with other SG components: the RBPs Pub1 and Pab1. Our experiments uncovered intimate relationships between protein evolution, protein dynamics and protein complex formation that exemplify various eIF4G1 (un)structure-function relationships. The N-terminus of eIF4G1 contains several short segments (BOXes) that are conserved in the *Saccharomycetales* family and act as short linear interaction motifs (SLiMs) for various purposes: (1) intramolecular and intermolecular self-recognition; (2) specific recognition of RBPs; and (3) RNA recognition ^28,29^.

The eIF4G1_1-249_ conformational ensemble is one of the few atomistic models of an IDP that, like other recent examples ^38,57-59^, reveals the importance of long-range interactions in these types of proteins. The eIF4G1_1-249_ ensemble shows a high degree of structural variability, as expected for an IDP, while maintaining a certain degree of compactness due to long-range contacts among several SLiMs. The prion-like BOX1 stands out among these eIF4G1_1-249_ motifs, participating in a fuzzy network of π-π and π-cation interactions with other motifs in the RNA1 domain and with BOX3. Arginine-aromatic contacts also have a π-π character due to the sp2 hybridization of the guanidinium group ^40^ and have been linked to the LLPS phenomenon ^8,39-41^. Our work shows that π-π contacts can also be important for stabilizing long IDRs and for masking prion-like elements such as BOX1. This unique element contains overlapping Pub1 binding (YNN) and amyloid-like (YYNN) motifs, two statistically underpopulated SLiMs in the yeast proteome although they are relatively abundant among Pub1 and eIF4G1 binding proteins and in SG proteins (Supplementary Figure 7). In nearly all these proteins the (YNN) and (YYNN) motifs are located in IDRs, suggesting that they might be involved in similar recognition events as those described here.

We found that Pub1 and Pab1 interact with various eIF4G1 elements as a double-key system (Figure 6C), promoting a reorganization of the conformational landscape of eIF4G1_1-249_ and triggering its oligomerization, likely through BOX1. These biophysical data could help to better understand the SG structure and dynamics and how these can be integrated in the stress signaling pathway (Figure 8). Eukaryotic cells have developed an integrated stress response pathway that senses different stimuli and triggers an enzymatic cascade that ends in the phosphorylation of eIF2-alpha ^60^. This important posttranslational modification inhibits translation by blocking the 48S pre-initiation complex (Figure 8). Pab1 and eIF4G1 are part of the 48S, together with other initiation factors and the ternary complex (eIF2-tRNAi-GTP). The eIF4G1 N-terminal IDR probably remains in a disordered stage, but stabilized by the intramolecular transient contacts that we described here, interactions with Pab1 (through BOX2 and BOX3), and perhaps transient contacts with the mRNA ^28^. We propose that the transition from the stalled 48S to the SG occurs by interaction between eIF4G1 IDR and Pub1 (and perhaps other RBPs). This would cause the remodeling of the transient contacts in the eIF4G1 IDR and unmask the amyloid-like properties of BOX1, facilitating the LLPS transition to SG in combination with other pro-amyloidogenic regions in Pab1 ^11^ and Pub1 ^12^.

Multivalence is a powerful driving force of phase transitions in proteins ^13^ and it is usually achieved through the concerted action of multiple elements. From this viewpoint, our study of the Pab1/Pub1/eIF4G1 system uncovers a set of new multivalent interactions that perhaps help to organize biological condensates at the molecular level, along the line of previous studies of the single components ^11,12^. These three proteins are abundant in cells, and therefore we propose that the interactions we characterized may describe some structural features of the SG cores ^6^. These substructures may develop early during the formation of SG and involve the oligomerization of individual mRNPs ^61^, perhaps guided by self-recognition of Pub1 and Pab1 as described here and previously ^52^.

Our model opens up new possibilities in the translation regulation field and reveal an intimate connection with SG biology through the central role of multivalent interactions between Pab1, Pub1 and eIF4G1 proteins.

## Acknowledgments

NMR experiments were performed in the “Manuel Rico” NMR laboratory (LMR) (CSIC). The synchrotron SAXS data was collected at beamline P12 operated by EMBL Hamburg at the PETRA III storage ring (DESY, Hamburg, Germany). The study was supported by funds from the Spanish MINECO (CTQ2018784371) to MJ and JMC-P, and Autonomous Community of Madrid (B2017/BMD73770) to JMP-C. BC-A and SM-L were funded by predoctoral grants from Spanish MINECO (BES-2015-073383) and Autonomous Community of Madrid (CPI/0265/2008) respectively. This work was supported by the Labex EpiGenMed, an « Investissements d’avenir » program (ANR-10-LABX-12-01) awarded to PB. The CBS is a member of France-BioImaging (FBI) and the French Infrastructure for Integrated Structural Biology (FRISBI), 2 national infrastructures supported by the French National Research Agency (ANR-10-INBS-04-01 and ANR-10-INBS-05, respectively). We would like to thank Clara M. Santiveri and Francisco Blanco for their helpful suggestions and critical reading of the manuscript and M.T. Seisdedos and G. Elvira (Confocal Laser and Multidimensional Microscopy Facility, CIB-CSIC) for assistance in imaging.

## Autor contributions

BC-A, SM-L, SC and JMP-C made all of the clones, mutants and recombinant proteins. BC-A, SM-L and JMP-C obtained and analyzed NMR data with the help of MJ. Residual Dipolar Couplings were measured by BC-A and NS, and analyzed with the help of PB and NS. SAXS experiments were performed by BC-A and PB and analyzed by SM-L and JMP-C with the advice of PB. Paramagnetic Relaxation Experiments were performed and analyzed by BC-A and JMP-C. Protein structure calculations were done by JMP-C. Confocal microscopy images were acquired and analyzed by BC-A, JMP-C and SZ with the help of SC. The project was conceived by JMP-C who wrote the paper with the assistance of BC-A and with contributions from SZ and other authors.

## MATERIAL AND METHODS

### Cloning, protein expression and purification

Plasmids and proteins used in this work are described in the key resources table. DNA fragments corresponding to wild-type constructs of eIF4G1, Pub1 and Pab1 were amplified from *Saccharomyces cerevisiae* genomic DNA using the DNA polymerases KOD or Pfu. These DNA fragments were cloned into a pET28-modified vector that contains an N-terminal thioredoxin A fusion tag, an internal 6xHis tag and a TEV protease site. eIF4G1 mutants were obtained using the Quick-change Lightning Kit and specific DNA primers. Plasmids corresponding to mutant and wild-type proteins were transformed into *E. coli* BL21 (DE3) competent cells and expressed in kanamycin-containing (30 µg/l) LB medium.

For isotope labelling of samples, a K-MOPS derived minimal medium ^62^ was supplemented with ^15^NH_4_Cl (1 g/l) and/or ^13^C-glucose (4 g/l). Cultures of eIF4G1 and its mutants were grown at 37 °C until OD_600nm_=0.6-0.8, when they were induced with 0.5 µM IPTG for 4 hours. Pab1 and Pub1 cultures, after reaching OD_600nm_=0.6, were transferred to 25 °C for induction with IPTG overnight (12-16 hours).

For purification of all recombinant proteins, cell pellets were resuspended in lysis buffer (25 mM potassium phosphate pH 8.0, 300 mM NaCl, 10 mM Imidazole and 1 tablet/50 ml of protease inhibitors cocktail), lysed by sonication and cleared by ultracentrifugation. The supernatant was purified by metal affinity chromatography using a HiTrap™ 5 ml column and elution with 25 mM potassium phosphate buffer pH 8.0, containing 300 mM NaCl and 300 mM imidazole. The samples containing the fusion protein were exchanged into 20 mM Tris pH 8.0 (in the case of the Pab1 construct this buffer was supplemented with 1 mM DTT), and digested overnight at 4 °C with homemade TEV protease. In the case of the Pub1 and Pab1 constructs, the samples were re-loaded onto the HiTrap nickel column to capture the protease, the cleaved N-terminal part of the fusion protein and the undigested protein. The flow through was further purified by ion exchange using an anion exchanger column (Q 5 ml) for all proteins except for Pub1 RRM3 that was purified with a cation exchange column (SP 5 ml). In either case, proteins were eluted with a linear salt gradient (to 1 M NaCl). In the case of the eIF4G1 construct, we observed that the second nickel column negatively affected protein stability and aggregation; we therefore purified the protein away from the uncleaved protein, thioredoxin A and TEV using a cation exchange column (SP 5 ml). Finally, the purified proteins were concentrated and the buffer was exchanged according to their intended use.

### Dynamic Light Scattering

The DLS measurements were carried out at 25 °C in a DynaPro Titan (Wyatt Technologies) instrument and were analysed with Dynamics V6 software. Protein mixtures (eIF4G1_1-249_, eIF4G1_1-249_ ΔBOX1, Pub1 RRM123 and Pab1 RRM12) were prepared from extensively centrifuged stocks (>1 h at 15000 RPM; 4 °C), and were filtered (0.22 µm) in PBS buffer and 200 g/l Ficoll 70 stock. Samples were mixed thoroughly and placed in a plastic cuvette (Eppendorf) for measurements. Individual correlation curves were recorded (10 acquisitions of 10s) every 15 minutes over a 5 h period.

### Small Angle X-ray Scattering (SAXS) measurements

SAXS experiments were performed using the P12-EMBL beamline at the DESY synchrotron in Hamburg and were analysed with ATSAS software. All data were collected in batch using 25 mM potassium phosphate pH 6.5 and 25 mM NaCl. The concentrations used for analysis of eIF4G1_1-249_ were 15 mg/ml, 10 mg/ml, 8 mg/ml, 5 mg/ml, 3 mg/ml and 1 mg/ml, those for Pub1 RRM123 were 12 mg/ml and 3mg/ml and those for SAXS Pab1 RRM12 experiments were 17 mg/ml, 10 mg/ml and 6 mg/ml. The final SAXS curves were generated using PRIMUS.

### NMR: resonance assignments and relaxation data

All samples were prepared in NMR buffer (25 mM potassium phosphate pH 6.5, 25 or 150 mM NaCl, 1 mM DTT and 10% D_2_O) and experimental data were acquired at 25 °C on a cryoprobe-equipped Bruker AV800 MHz spectrometer. Assignment of the backbone ^1^H, ^15^N and ^13^C atoms was achieved by following the standard methodology. The 3D HNCA, HNCO, HN(CO)CA, CBCA(CO)NH and CBCANH experiments were used for backbone assignment and 3D (H)CCH-TOCSY were recorded to assign side chain resonances (^63^ and the references therein). Protein concentrations ranged between 100-200 µM. The chemical shifts were deposited in the Biomagnetic Resonance Database (BMRB) with codes 28121 (eIF4G1_1-249_) and 34517 (eIF4G1_35-49_-Pub1 RRM3). The ^15^N backbone amide relaxation T_1_ and T_2_ parameters were measured with series of ^1^H-^15^N spectra of standard inversion-recovery and Carr-Purcell-Meiboom-Gill sequences (CPMG). NMR spectra were processed using TOPSPIN v2.1 (Bruker) and NMRPipe, and analyses were done with CcpNmr Analysis.

### NMR: Residual dipolar couplings and PRE measurements

The filamentous phage Pf1 was used at a final concentration of 20 mg/ml to induce weak alignment of eIF4G1_1-249_ (200 µM in 25 mM potassium phosphate pH 6.5 and 25 mM NaCl).The NMR experiments were carried out at 298 K in a Bruker Avance III 800 MHz spectrometer equipped with a cryogenic triple resonance probe. Two samples (isotropic and anisotropic) were prepared and couplings (J and J+D) were measured with ^15^N-HSQC-DSSE (In Phase Anti Phase IPAP). Experiments were processed using TOPSPIN v2.1 and NMRPipe, and were analyzed with the program CcpNmr Analysis.

For the paramagnetic relaxation enhancement, protein samples from different cysteine-containing eIF4G1_1-249_ mutants (S200C and Q109C) were chemically modified using the following protocol. Mutant protein samples (600-700 µM) were pre-treated with 5 mM DTT for two hours at room temperature. The DTT was then eliminated by fast buffer exchange into 25 mM Tris pH 9.0 and 25 mM NaCl using a Nap-5 desalting column. Labelling with 4-(2-Iodoacetamido)-TEMPO was initiated immediately after column elution by adding a tenfold molar excess of the probe dissolved in ethanol (25 mM spin label stock). The reaction was allowed to proceed for 30 minutes at room temperature in the dark. The excess iodoacetamide label was quenched with 10 mM 2-mercaptoethanol for 10 minutes, and afterwards the protein adduct was exchange into 25 mM potassium phosphate pH 6.5, 25 mM NaCl and 1 mM DTT for later use. The NMR samples were prepared in 5 mm tubes sealed in a N_2_ atmosphere to avoid reduction by air, and high resolution ^1^H-^15^N HSQC spectra were recorded for the oxidized state (active spin label). Subsequently, the spin label was reduced with 10 µM ascorbate ^64^, and the reference ^1^H-^15^N HSQC without paramagnetic relaxation enhancement was recorded. The relaxation effect was calculated as the intensity ratios between peaks in the two spectra.

### Structure calculations of eIF4G1_35-49_-Pub1 RRM3 and the eIF4G BOX3 peptide

The NMR structure of the eIF4G_187-234_ construct was determined from NOE-derived distance restraints (2D NOESY spectrum with 60 <u>ms</u> mixing time) and angular restraints (from 13C chemical shifts and TALOS+) using the program Cyana. Protein assignments were obtained by comparison with other eIF4G1 constructs and confirmed by triple resonance 3D spectra: CBCA(CO)NH, HNCACB and HNCO.

Two different chimeras of Pub1 with eIF4G1_37-51_ were constructed, with eIF4G1_35-49_ fused either to the N- or C-terminus of Pub1 RRM3. Of these constructs, only the N-terminal fusion (eIF4G1_35-49_) proved to have the right topology and the structure was determined using a similar protocol to analysis of the structure of Pub1 RRM3 ^65^ using distance restraints from a 2D NOESY (60 ms mixing time).

### Structure calculation of eIF4G1_**1-249**_

eIF4G1_1-249_ structures (80000) were calculated with the program Cyana 3.0 using: 1) experimental NOE-derived distance restraints for the BOX3 region; 2) π-π interactions between Tyr, Phe and Trp; and 3) π-cation interactions between Arg and Tyr/Phe/Trp. The latter two interaction types were referred to as knowledge-based constraints (K-BC) and were included, given the importance of these types of contacts for IDP interactions (see main text for references). For each individual structure calculation, the origin residue (Arg/Tyr/Phe/Trp) was randomly selected (80% probability) and ambiguous restraints were generated to the other interaction partner (Arg/Tyr/Phe/Trp). In this way, each of the individual structure calculations contains a unique set of knowledge-based distance restraints. This protocol ensures high variability by avoiding biases of specific pairwise iterations. Thus the final interactions present on each conformer are freely selected during the calculations. A similar protocol was followed to calculate the structures (80000) of eIF4G1_1-249_ dimers, using the same experimental restraints and ambiguous Tyr-Tyr contacts as dimerization driving interactions.

The theoretical PRE-derived intensity ratios were calculated using equations in ^37^ (Supplementary Figure 5) for the eIF4G1_1-249_ monomers and dimers. Correlation time τ_c_ was estimated from the averaged T_1_/T_2_ and *d* was estimated from the distances between amide backbones (N) and Q109/S200 side-chains (Cβ). We next used a home-made greedy algorithm to select the ensembles that better reproduced the PRE profiles. The algorithm calculates the residual to the experimental data (Q109 and S200) and chooses, as a seed, the conformer that better agrees (lower sum of residuals) with the data. In the next steps, the residuals were computed across 2, 3,.., n structures always choosing the combination with minimal violations. The procedure was repeated until the ensemble size reached 2000 conformers (∼2.5% of the original). The first 500 structures were included in the final eIF4G1_1-249_ monomer and eIF4G1_1-249_ dimer ensembles (PRE-derived).

We used the EOM protocol to fit the SAXS data. Pools of theoretical SAXS curves were constructed for the eIF4G1_1-249_ monomers (2000 conformers) and dimers (2000 conformers). These two pools were combined with the genetic algorithm in the EOM program to model the curve with a fixed size ensemble (50 structures). The percentages of each pool were freely selected by the algorithm and the procedure. The procedure was repeated 10 times, obtaining a 500-member set. It should be noted that some of the structures are repeated between individual EOM calculations. The theoretical values of the PREs, and other structural properties, were calculated as averages across the different ensembles using home-made perl scripts.

### Confocal microscopy

eIF4G1_1-249_, Pab1 RRM12 and Pub1 RRM123 proteins were purified as described above and were labelled with Alexa Fluor 488 or Alexa Fluor 647 carboxylic acid succinimidyl ester dyes (Molecular Probes), using protein to probe ratios of 1:3. The coupling reaction was carried out in the dark in PBS (pH 7.4) buffer for 30 minutes on ice and the unreacted probe was removed by size exclusion chromatography using a Nap-5 column. Samples for visualization were prepared by mixing eIF4G1_1-249_ with Pub1 RRM123 and Pab1 RRM12 in different combinations and at concentrations of 1, 5 and 20 µM respectively, in PBS, 0.1 mM DTT (pH 7.4) buffer and 200 g/l Ficoll 70. Samples, that contain 1 µM of fluorescently labelled protein (Alexa Fluor 488-labelled eIF4G1 and/or Alexa Fluor 647-labelled Pub1 or Pab1) for visualization, were placed in silicone chambers that were glued to coverslips and were visualized with Leica TCS SP2 or TCS-SP5 inverted confocal microscopes with a HCX PL APO 63x oil immersion objective (N.A. = 1.4; Leica, Mannheim, Germany). Alexa 488 and Alexa 467 were excited using 488 nm and 633 nm laser excitation lines, respectively. The concentration of the various Alexa-labelled proteins was kept at 1 µM and the solution was supplemented with unlabeled proteins to reach the above-mentioned concentrations. Various images were registered for each sample, corresponding to different observation fields.

## REFERENCES

1. Crawford, R. & Pavitt, G. Translational regulation in response to stress in Saccharomyces cerevisiae. Yeast 36, 5–21 (2019).

2. Kedersha, N.L., Gupta, M., Li, W., Miller, I. & Anderson, P. RNA-binding proteins TIA-1 and TIAR link the phosphorylation of eIF-2 alpha to the assembly of mammalian stress granules. J Cell Biol 147, 1431–42 (1999).

3. Hoyle, N.P., Castelli, L.M., Campbell, S.G., Holmes, L.E. & Ashe, M.P. Stress-dependent relocalization of translationally primed mRNPs to cytoplasmic granules that are kinetically and spatially distinct from P-bodies. J Cell Biol 179, 65–74 (2007).

4. Buchan, J., Muhlrad, D. & Parker, R. P bodies promote stress granule assembly in Saccharomyces cerevisiae. Journal of Cell Biology 183, 441–455 (2008).

5. Buchan, J., Yoon, J. & Parker, R. Stress-specific composition, assembly and kinetics of stress granules in Saccharomyces cerevisiae. Journal of Cell Science 124, 228–239 (2011).

6. Jain, S. et al. ATPase-Modulated Stress Granules Contain a Diverse Proteome and Substructure. Cell 164, 487–98 (2016).

7. Shin, Y. & Brangwynne, C.P. Liquid phase condensation in cell physiology and disease. Science 357(2017).

8. Brangwynne, C., Tompa, P. & Pappu, R. Polymer physics of intracellular phase transitions. Nature Physics 11, 899–904 (2015).

9. Nakashima, K.K., Vibhute, M.A. & Spruijt, E. Biomolecular Chemistry in Liquid Phase Separated Compartments. Front Mol Biosci 6, 21 (2019).

10. Lin, Y., Protter, D.S., Rosen, M.K. & Parker, R. Formation and Maturation of Phase-Separated Liquid Droplets by RNA-Binding Proteins. Mol Cell 60, 208–19 (2015).

11. Riback, J.A. et al. Stress-Triggered Phase Separation Is an Adaptive, Evolutionarily Tuned Response. Cell 168, 1028-1040.e19 (2017).

12. Kroschwald, S. et al. Different Material States of Pub1 Condensates Define Distinct Modes of Stress Adaptation and Recovery. Cell Rep 23, 3327–3339 (2018).

13. Li, P. et al. Phase transitions in the assembly of multivalent signalling proteins. Nature 483, 336–40 (2012).

14. Banani, S.F., Lee, H.O., Hyman, A.A. & Rosen, M.K. Biomolecular condensates: organizers of cellular biochemistry. Nat Rev Mol Cell Biol 18, 285–298 (2017).

15. Buchan, J. & Parker, R. Eukaryotic Stress Granules: The Ins and Outs of Translation. Molecular Cell 36, 932–941 (2009).

16. Protter, D.S.W. & Parker, R. Principles and Properties of Stress Granules. Trends Cell Biol 26, 668–679 (2016).

17. Guzikowski, A.R., Chen, Y.S. & Zid, B.M. Stress-induced mRNP granules: Form and function of processing bodies and stress granules. Wiley Interdiscip Rev RNA 10, e1524 (2019).

18. Kedersha, N. & Anderson, P. Regulation of translation by stress granules and processing bodies. Prog Mol Biol Transl Sci 90, 155–85 (2009).

19. Sonenberg, N. & Hinnebusch, A.G. Regulation of translation initiation in eukaryotes: mechanisms and biological targets. Cell 136, 731–45 (2009).

20. Preiss, T. & Hentze, M.W. From factors to mechanisms: translation and translational control in eukaryotes. Curr Opin Genet Dev 9, 515–21 (1999).

21. Aitken, C.E. & Lorsch, J.R. A mechanistic overview of translation initiation in eukaryotes. Nat Struct Mol Biol 19, 568–76 (2012).

22. Wells, S.E., Hillner, P.E., Vale, R.D. & Sachs, A.B. Circularization of mRNA by eukaryotic translation initiation factors. Mol Cell 2, 135–40 (1998).

23. Tarun, S.Z., Wells, S.E., Deardorff, J.A. & Sachs, A.B. Translation initiation factor eIF4G mediates in vitro poly(A) tail-dependent translation. Proc Natl Acad Sci U S A 94, 9046–51 (1997).

24. Goyer, C. et al. TIF4631 and TIF4632: two yeast genes encoding the high-molecular-weight subunits of the cap-binding protein complex (eukaryotic initiation factor 4F) contain an RNA recognition motif-like sequence and carry out an essential function. Mol Cell Biol 13, 4860–74 (1993).

25. Tarun, S.Z. & Sachs, A.B. Association of the yeast poly(A) tail binding protein with translation initiation factor eIF-4G. EMBO J 15, 7168–77 (1996).

26. Kessler, S.H. & Sachs, A.B. RNA recognition motif 2 of yeast Pab1p is required for its functional interaction with eukaryotic translation initiation factor 4G. Mol Cell Biol 18, 51–7 (1998).

27. Santiveri, C.M., Mirassou, Y., Rico-Lastres, P., Martínez-Lumbreras, S. & Pérez-Cañadillas, J.M. Pub1p C-Terminal RRM Domain Interacts with Tif4631p through a Conserved Region Neighbouring the Pab1p Binding Site. Plos One 6(2011).

28. Park, E.H. et al. Multiple elements in the eIF4G1 N-terminus promote assembly of eIF4G1 PABP mRNPs in vivo. EMBO J 30, 302–16 (2011).

29. Berset, C., Zurbriggen, A., Djafarzadeh, S., Altmann, M. & Trachsel, H. RNA-binding activity of translation initiation factor eIF4G1 from Saccharomyces cerevisiae. RNA 9, 871–80 (2003).

30. Schütz, P. et al. Crystal structure of the yeast eIF4A-eIF4G complex: an RNA-helicase controlled by protein-protein interactions. Proc Natl Acad Sci U S A 105, 9564–9 (2008).

31. Hershey, P.E. et al. The Cap-binding protein eIF4E promotes folding of a functional domain of yeast translation initiation factor eIF4G1. J Biol Chem 274, 21297–304 (1999).

32. Poornima, G., Shah, S., Vignesh, V., Parker, R. & Rajyaguru, P.I. Arginine methylation promotes translation repression activity of eIF4G-binding protein, Scd6. Nucleic Acids Res 44, 9358–9368 (2016).

33. Gibbs, E., Cook, E. & Showalter, S. Application of NMR to studies of intrinsically disordered proteins. Archives of Biochemistry and Biophysics 628, 57–70 (2017).

34. Milles, S., Salvi, N., Blackledge, M. & Jensen, M.R. Characterization of intrinsically disordered proteins and their dynamic complexes: From in vitro to cell-like environments. Prog Nucl Magn Reson Spectrosc 109, 79–100 (2018).

35. Robinson, N.E. & Robinson, A.B. Prediction of protein deamidation rates from primary and three-dimensional structure. Proc Natl Acad Sci U S A 98, 4367–72 (2001).

36. Camilloni, C., De Simone, A., Vranken, W. & Vendruscolo, M. Determination of Secondary Structure Populations in Disordered States of Proteins Using Nuclear Magnetic Resonance Chemical Shifts. Biochemistry 51, 2224–2231 (2012).

37. Battiste, J.L. & Wagner, G. Utilization of site-directed spin labeling and high-resolution heteronuclear nuclear magnetic resonance for global fold determination of large proteins with limited nuclear overhauser effect data. Biochemistry 39, 5355–65 (2000).

38. Schwalbe, M. et al. Predictive atomic resolution descriptions of intrinsically disordered hTau40 and α-synuclein in solution from NMR and small angle scattering. Structure 22, 238–49 (2014).

39. Wang, J. et al. A Molecular Grammar Governing the Driving Forces for Phase Separation of Prion-like RNA Binding Proteins. Cell 174, 688-699.e16 (2018).

40. Vernon, R.M. et al. Pi-Pi contacts are an overlooked protein feature relevant to phase separation. Elife 7(2018).

41. Gomes, E. & Shorter, J. The molecular language of membraneless organelles. Journal of Biological Chemistry 294, 7115–7127 (2019).

42. Bhattacharya, S. & Lin, X. Recent Advances in Computational Protocols Addressing Intrinsically Disordered Proteins. Biomolecules 9(2019).

43. Kragelj, J., Blackledge, M. & Jensen, M.R. Ensemble Calculation for Intrinsically Disordered Proteins Using NMR Parameters. Adv Exp Med Biol 870, 123–47 (2015).

44. Kurzbach, D., Kontaxis, G., Coudevylle, N. & Konrat, R. NMR Spectroscopic Studies of the Conformational Ensembles of Intrinsically Disordered Proteins. Adv Exp Med Biol 870, 149–85 (2015).

45. Das, P., Matysiak, S. & Mittal, J. Looking at the Disordered Proteins through the Computational Microscope. Acs Central Science 4, 534–542 (2018).

46. Bernadó, P. et al. A structural model for unfolded proteins from residual dipolar couplings and small-angle x-ray scattering. Proc Natl Acad Sci U S A 102, 17002–7 (2005).

47. Estaña, A. et al. Realistic Ensemble Models of Intrinsically Disordered Proteins Using a Structure-Encoding Coil Database. Structure 27, 381-391.e2 (2019).

48. Güntert, P. & Buchner, L. Combined automated NOE assignment and structure calculation with CYANA. J Biomol NMR 62, 453–71 (2015).

49. Bernadó, P., Mylonas, E., Petoukhov, M.V., Blackledge, M. & Svergun, D.I. Structural characterization of flexible proteins using small-angle X-ray scattering. J Am Chem Soc 129, 5656–64 (2007).

50. Tria, G., Mertens, H.D., Kachala, M. & Svergun, D.I. Advanced ensemble modelling of flexible macromolecules using X-ray solution scattering. IUCrJ 2, 207–17 (2015).

51. Hajizadeh, N.R., Franke, D., Jeffries, C.M. & Svergun, D.I. Consensus Bayesian assessment of protein molecular mass from solution X-ray scattering data. Sci Rep 8, 7204 (2018).

52. Yao, G. et al. PAB1 self-association precludes its binding to poly(A), thereby accelerating CCR4 deadenylation in vivo. Mol Cell Biol 27, 6243–53 (2007).

53. Schäfer, I.B. et al. Molecular Basis for poly(A) RNP Architecture and Recognition by the Pan2-Pan3 Deadenylase. Cell 177, 1619-1631.e21 (2019).

54. Deo, R., Bonanno, J., Sonenberg, N. & Burley, S. Recognition of polyadenylate RNA by the poly(A)-binding protein. Cell 98, 835–845 (1999).

55. Safaee, N. et al. Interdomain Allostery Promotes Assembly of the Poly(A) mRNA Complex with PABP and eIF4G. Molecular Cell 48, 375–386 (2012).

56. Cohen, R.D. & Pielak, G.J. A cell is more than the sum of its (dilute) parts: A brief history of quinary structure. Protein Sci 26, 403–413 (2017).

57. Kubáň, V. et al. Quantitative Conformational Analysis of Functionally Important Electrostatic Interactions in the Intrinsically Disordered Region of Delta Subunit of Bacterial RNA Polymerase. J Am Chem Soc 141, 16817–16828 (2019).

58. Shrestha, U.R. et al. Generation of the configurational ensemble of an intrinsically disordered protein from unbiased molecular dynamics simulation. Proc Natl Acad Sci U S A 116, 20446–20452 (2019).

59. Cordeiro, T.N. et al. Interplay of Protein Disorder in Retinoic Acid Receptor Heterodimer and Its Corepressor Regulates Gene Expression. Structure 27, 1270-1285.e6 (2019).

60. Pakos-Zebrucka, K. et al. The integrated stress response. EMBO Rep 17, 1374–1395 (2016).

61. Wheeler, J.R., Matheny, T., Jain, S., Abrisch, R. & Parker, R. Distinct stages in stress granule assembly and disassembly. Elife 5(2016).

62. Neidhardt, F.C., Bloch, P.L. & Smith, D.F. Culture Medium for Enterobacteria. Journal of Bacteriology 119, 736–747 (1974).

63. Sattler, M., Schleucher, J. & Griesinger, C. Heteronuclear multidimensional NMR experiments for the structure determination of proteins in solution employing pulsed field gradients. Progress in Nuclear Magnetic Resonance Spectroscopy 34, 93–158 (1999).

64. Gillespie, J.R. & Shortle, D. Characterization of long-range structure in the denatured state of staphylococcal nuclease .1. Paramagnetic relaxation enhancement by nitroxide spin labels. Journal of Molecular Biology 268, 158–169 (1997).

65. Santiveri, C.M., Mirassou, Y., Rico-Lastres, P., Martinez-Lumbreras, S. & Perez-Canadillas, J.M. Pub1p C-terminal RRM domain interacts with Tif4631p through a conserved region neighbouring the Pab1p binding site. PLoS One 6, e24481 (2011).

