## Supplementary for "Multivalent interactions between eIF4G1, Pub1 and Pab1 drive the formation of protein condensates"

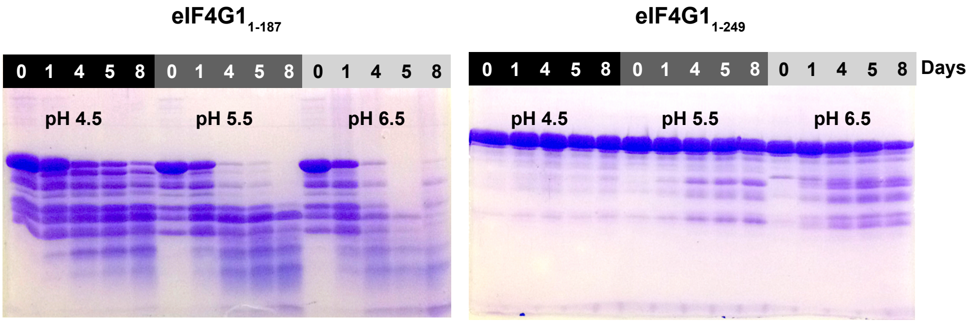


**Supplementary Figure 1.** Stability of eIF4G1$_{1\text{-}187}$ and eIF4G1$_{1\text{-}249}$ constructs at equivalent concentrations (50 µM) and under different buffer conditions. Samples were incubated at 25 °C for 0, 1, 4, 5, or 8 days in 25 mM sodium acetate (pH 4.5, 5.5) or potassium phosphate buffer (pH 6.5) containing 150 mM NaCl and 0.0005% $NaN_3$. Samples were then electrophoresed and the gels were labelled with Coomassie Blue. The construct lacking the BOX3 element (eIF4G1$_{1\text{-}187}$) is less stable than that with this element (eIF4G1$_{1\text{-}187}$).

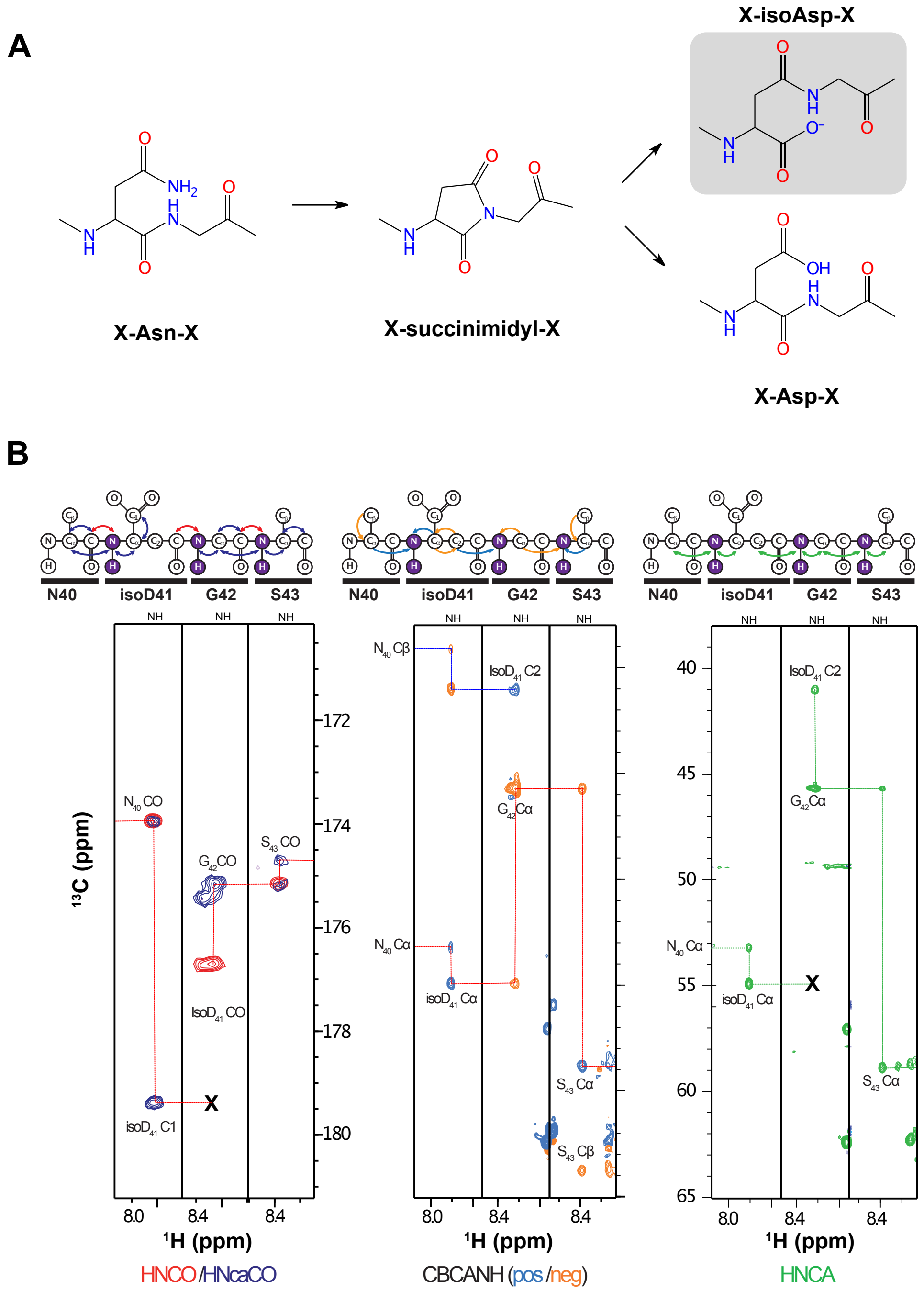


**Supplementary Figure 2. (A)** Schematic view of the chemical process of deamidation of asparagines. The pentacyclic intermediate could evolve to isoD or D, but only the first pathway was detected for eIF4G1$_{1\text{-}249}$. **(B)** $^{15}$N planes of various triple resonance experiments showing crosspeaks within the isoD$_{41}$-G$_{42}$-S$_{43}$ segment indicating the sequential connectivities. The expected magnetization transfers for each spectrum type are shown above with arrows colored in the same color as the experimental data. The sequential connections are interrupted in HNCA, HNCO and HNCACO due to the β-configuration of the backbone at isoD41.

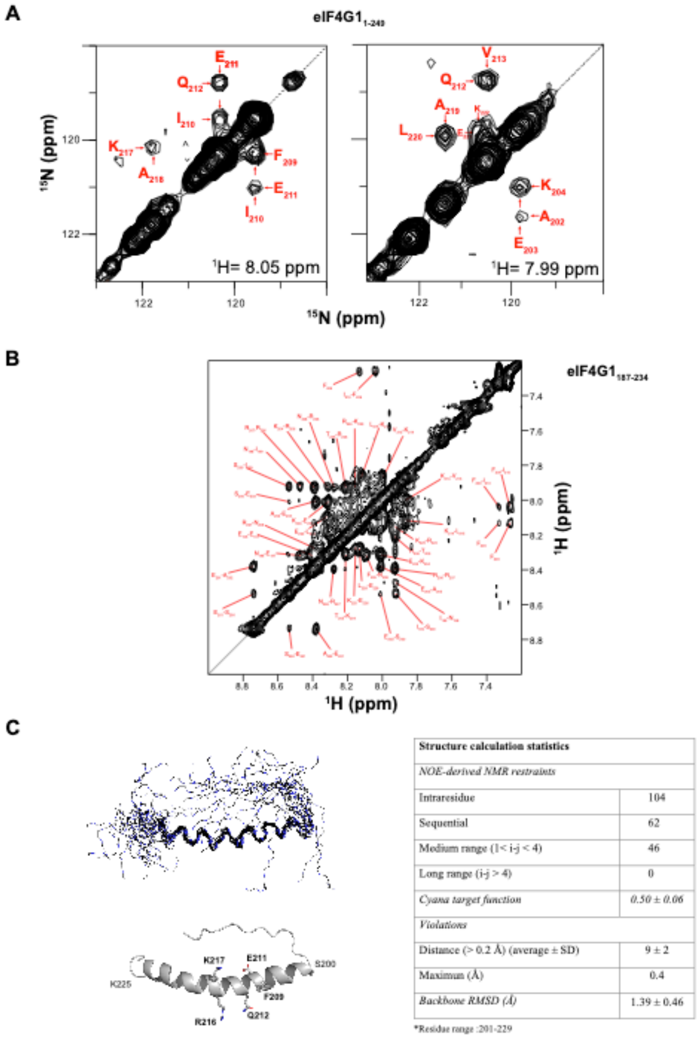


| Structure calculation statistics | |
|---|---|
| *NOE-derived NMR restraints* | |
| Intraresidue | 104 |
| Sequential | 62 |
| Medium range (1< i-j < 4) | 46 |
| Long range (i-j > 4) | 0 |
| *Cyana target function* | *0.50 ± 0.06* |
| *Violations* | |
| Distance (> 0.2 Å) (average ± SD) | 9 ± 2 |
| Maximum (Å) | 0.4 |
| *Backbone RMSD (Å)* | 1.39 ± 0.46 |

*Residue range :201-229

**Supplementary Figure 3. (A)** The $^{15}N$-$^{15}N$ planes in the 3D $^{1}H$-$^{15}N$-HSQC-NOESY-$^{1}H$ $^{15}N$-HSQC spectra of eIF4G1$_{1-249}$ show the characteristic sequential HN-HN NOEs between consecutive residues of the α-helix in BOX3. **(B)** Detail of the HN-HN NOEs in the 2D NOESY of eIF4G1$_{187-234}$. **(C)** The NMR structure of this eIF4G1$_{187-234}$ construct forms a continuous, slightly curved, α-helix with 7-turns in which the conserved residues

(F209, E211, Q212, R216 and K217) are located in the middle. Structure calculation statistics are shown on the right.

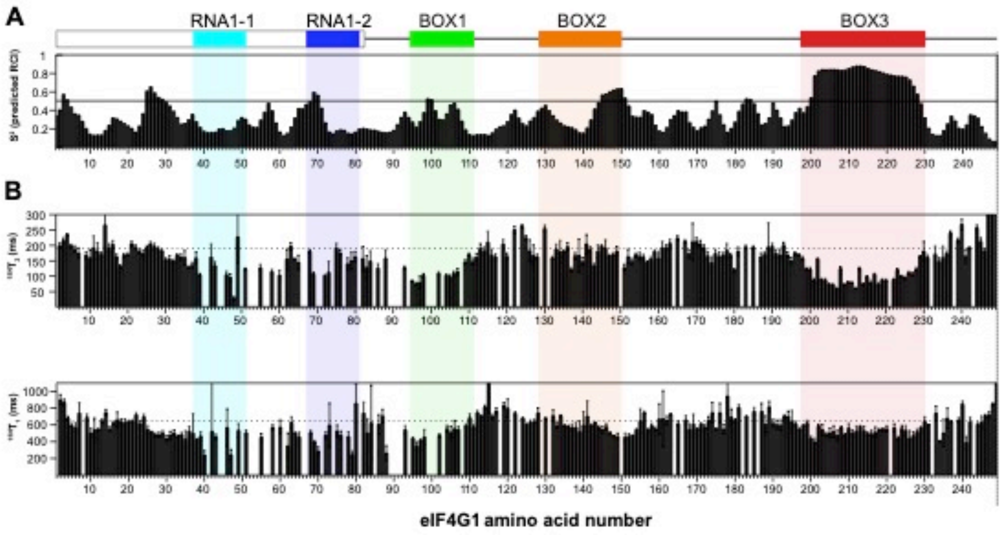


**Supplementary Figure 4**. **(A)** Random coil index (RCI) S2 values predicted from chemical shifts of eIF4G1$_{1\text{-}249}$ using the program Camshift. **(B)** Experimental values of $^{15}$N $T_1$ and $T_2$ at 25 ºC in an Bruker AV800 spectrometer. Amino acids located in eIF4G1$_{1\text{-}249}$ conserved regions are indicated in color.

**Eq. 1** $$\frac{I_{ox}}{I_{red}} = \frac{R_2 e^{-(R_2^{spin} t)}}{R_2 + R_2^{spin}}$$

**Eq. 2** $$R_2^{spin} = \left[\frac{K}{d^6}\left(4\tau_c + \frac{3\tau_c}{1+\omega_I^2\tau_c^2}\right)\right]$$

**Eq. 3** $$R_2 = \pi\Delta\nu_{1/2}$$

**Eq. 4** $$\tau_c = \frac{1}{4\pi\Delta\nu_N}\sqrt{6\frac{T_1^N}{T_2^N} - 7}$$

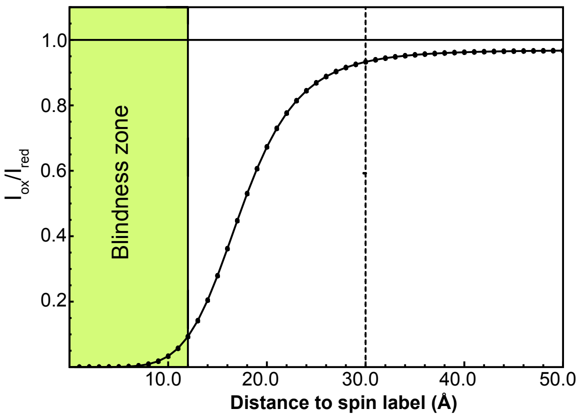


**Supplementary Figure 5.** The theoretical intensity ratios of oxidized (ox) to reduced (red) forms of the spin-labeled eIF4G1$_{1\text{-}249}$ mutants were computed as previously described in (Battiste and Wagner, 2000) using eq. 1 and 2. Proton $R_2$ relaxation was estimated from the average of cross peak linewidths at half-height (eq. 3) and correlation time ($\tau_c$) from the average $T^N_1/T^N_2$ values (Figure 2A middle panel). The functional dependence of eq.1 on distance is shown in the graph on the right. The vertical line at 30 Å marks the Paramagnetic Relaxation Enhancement (PRE) upper limit used for all the experimental $I_{ox}/I_{red}$ values below 0.8.

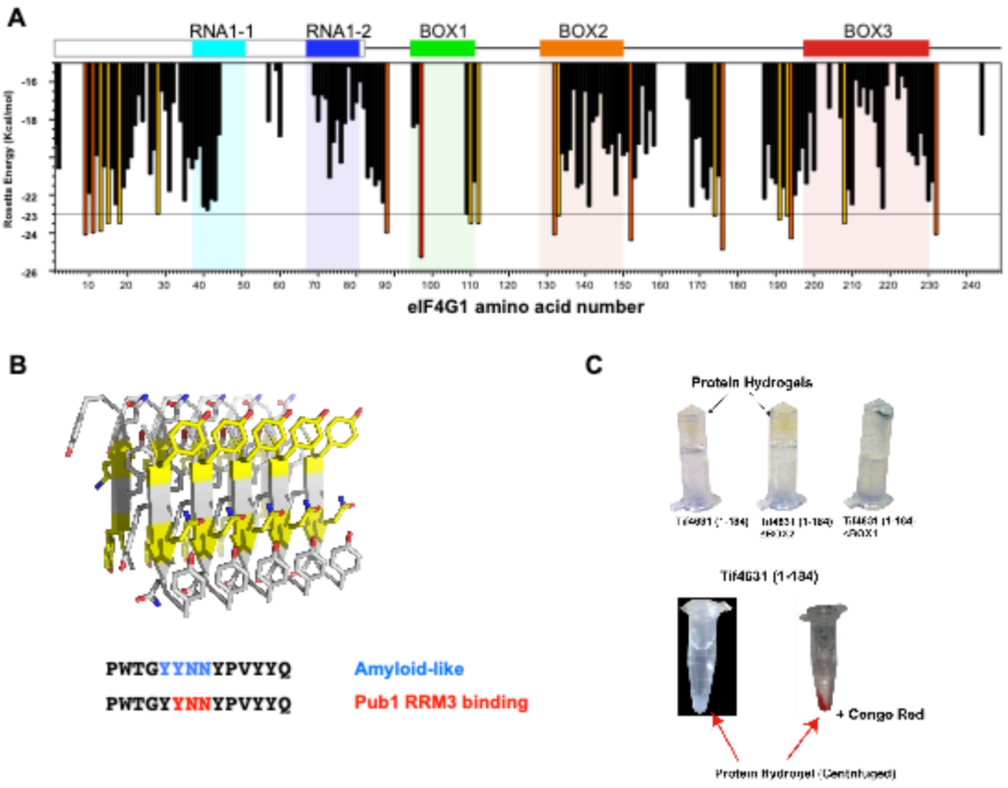


D

| Protein | Motifs | eIF4G | Pub1 | SG core | Observations |
|---|---|---|---|---|---|
| ARP3 | RKGLYNNIVLS | X | | | |
| BIO3 | VIHGYNNPELN | X | | X | Metabolite biosynthesis |
| CYS4 | ILDQYNNMMNP | X | | X | Metabolite biosynthesis |
| DED1 | NSSNYNNNNGG | X | | X | RNA helicase |
| FAB1 | LFAFYNNRKKD<br>NNSNYNNTNSN | | | | Kinase |
| KSP1 | DDEYYNNLDEG<br>QNRNYNNNNNN<br>HGSNYNNFNNG | X | | X | Kinase |
| LEU1 | GDIFYNNSFKN | X | | X | Metabolite biosynthesis |
| MCM7 | LPVDYNNLFNE | | | | DNA helicase |
| MRN1 | MVVSYNNNNNN | X | | X | RBP: 5 RRMs |
| NAB6 | CLNFYNNILQR | X | X | X | RBP:3 RRM, Nab6_mRNP_bdg |
| NGR1 | NYYNYNNYDYH | | X | X | RBP: 3 RRM |
| NRP1 | PNYRYNNNINN | | X | X | RBP: RRM,2 ZF |
| PBS2 | TSSHYNNINAD | | | X | Kinase |
| SLF1 | KRGKYNNDINK<br>GKTRYNNNSRH<br>QQQYYNNINYQ | X | | X | RBP: HTH-Latype |
| SYT1 | PTSVYNNKSNA | X | | | |
| TIF4631 | GYTNYNNGSNY<br>GPNRYNNRGNY<br>WTGYYNNYPVY | X | X | X | eIF4G1 |
| TIF4632 | GTNKYNNRNNS<br>NAQYYNNRFNN | X | X | X | eIF4G2 |
| XPT1 | MYISYNNIHKL | X | | | Metabolite biosynthesis |
| RIE1 | QHKVYNNHNSH<br>PMYDYNNYTYD | | | X | RBP: 5 RRMs |

**Supplementary Figure 6. (A)** Fibrillation propensity of hexapeptide sequences in eIF4G1$_{1\text{-}249}$ calculated using ZIPPERDB https://services.mbi.ucla.edu/zipperdb/. Segments whose Rosetta models have energies below -23 Kcal/mol are coloured in yellow (-23.0 to -24. Kcal/mol), orange (-24.0 to -25.0 Kcal/mol) and red (< -25.0 Kcal/mol). See program reference for specific details. **(B)**. Amyloid model for eIF4G1$_{96\text{-}102}$. **(C)** Hydrogel forming capacity of aged samples of different eIF4G1 constructs. The hydrogels were stained with amyloid-binding Congo-Red. **(D)** Distribution of the YYNN and YNN motifs in the S. cerevisiae proteome. Columns show the sequences alignments, presence on eIF4G, in Pub1 interactomes (from the SGD database) and in the stress granule (SG) core particles (Jain et al., 2016).

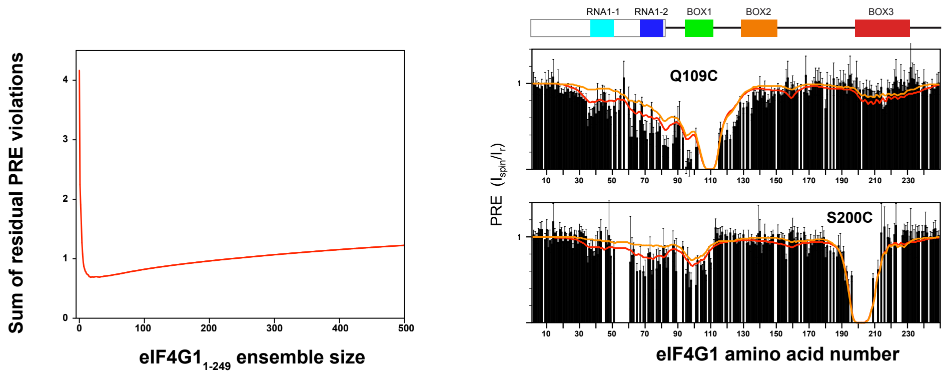


**Supplementary Figure 7.** The left panel shows the evolution of Residual PRE violations (as defined in materials and methods) as a function of the eIF4G1$_{1\text{-}249}$ ensemble size. The right panel shows the comparison of the calculated PRE values from the PRE-selected (red) and the EOM selected ensembles (orange). See materials and methods for further details.

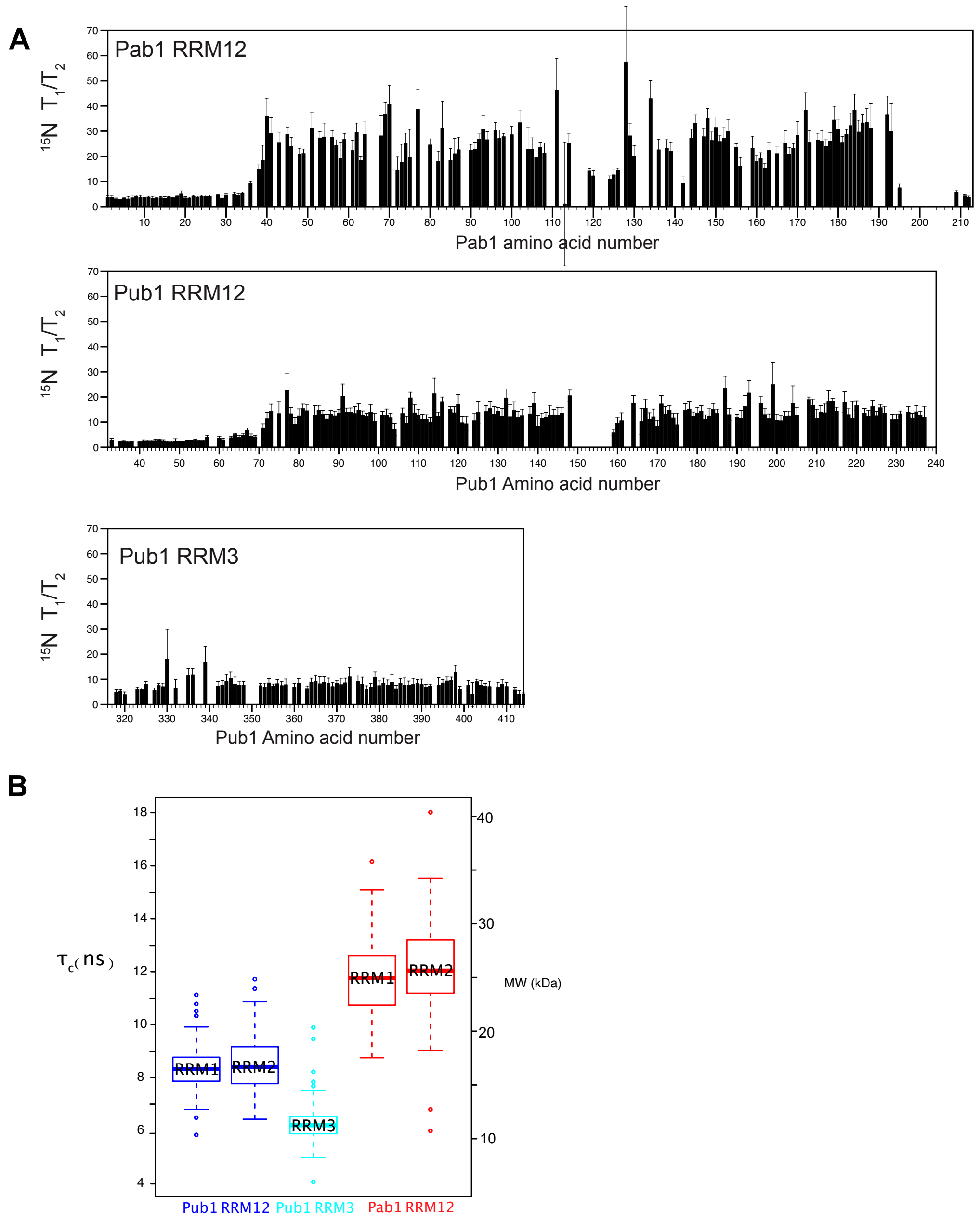


**Supplementary Figure 8. (A)** $^{15}$N relaxation analysis of Pub1 and Pab1 constructs. All the proteins were analyzed using the same concentration and buffer conditions. The experiments were recorded in a Bruker AV800 magnet at 25 ºC and analyzed with ccpnmrAnalysis software. **(B)** Correlation times ($\tau_c$) computed from the subsets of data in A delimited for the different RRM boundaries. Upper and lower box limits delimited Q1 and Q3 quartiles and the horizontal bar the median. The whiskers delimited the range excluding the out layers (dots). The scale on the right represent the equivalent molecular weight for a spherical protein model (a good approximation of the RRM shape).

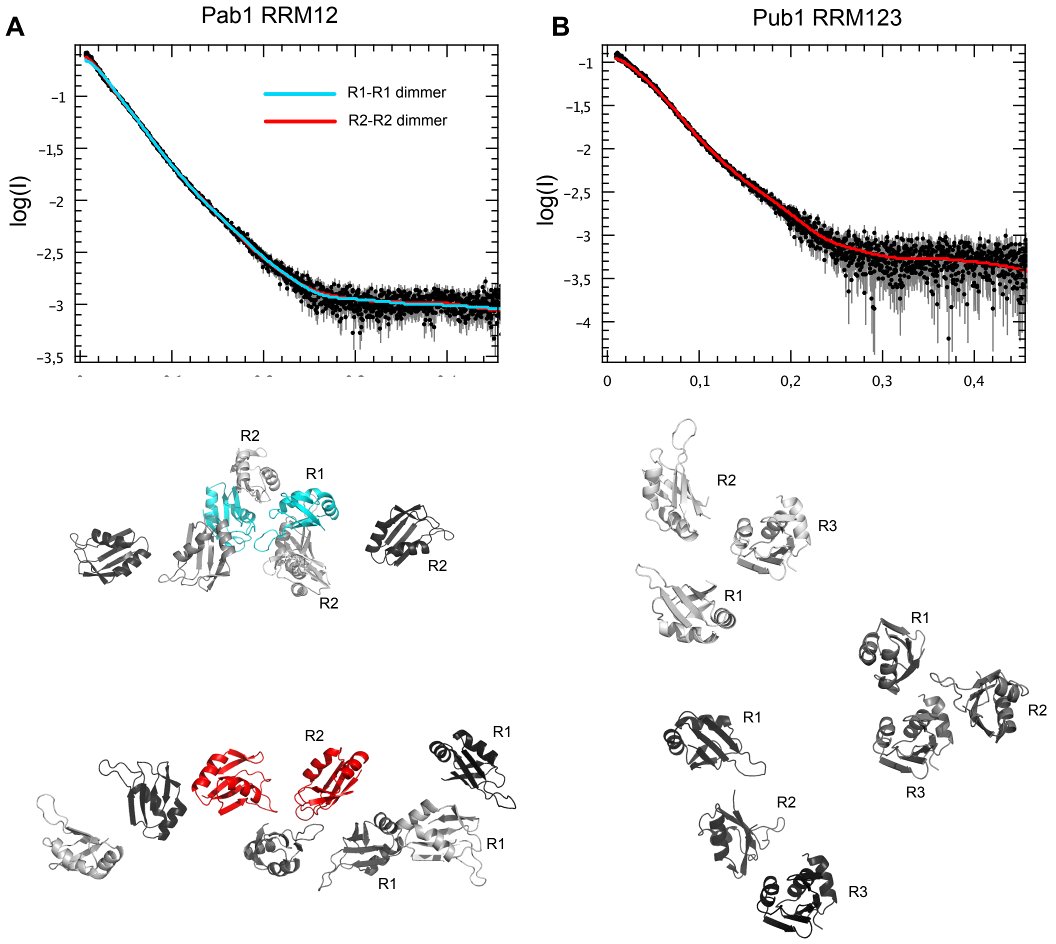


**Supplementary Figure 9**. EOM calculations for Pab1 RRM12 **(A)** and Pub1 RRM123 **(B)**. Fitting to the experimental curves and some examples of the modeled structures are shown. The conformers of Pab1 RRM1 were structurally aligned by either RRM1 (cyan) or RRM2 (red) for EOM calculations based on RRM1 or RRM2 dimerization. Other RRMs are indicated in different shades of gray depending on the individual conformer they belong to. The corresponding fitting curves are shown above as cyan (RRM1 dimers) and red lines (RRM2 dimers). Conformers of Pub1 RRM123 monomers calculated by the EOM calculations are shown below the SAXS data **(B)**. Individual RRMs of Pub1 are labelled. The corresponding fitting curve is shown in red in the graph.

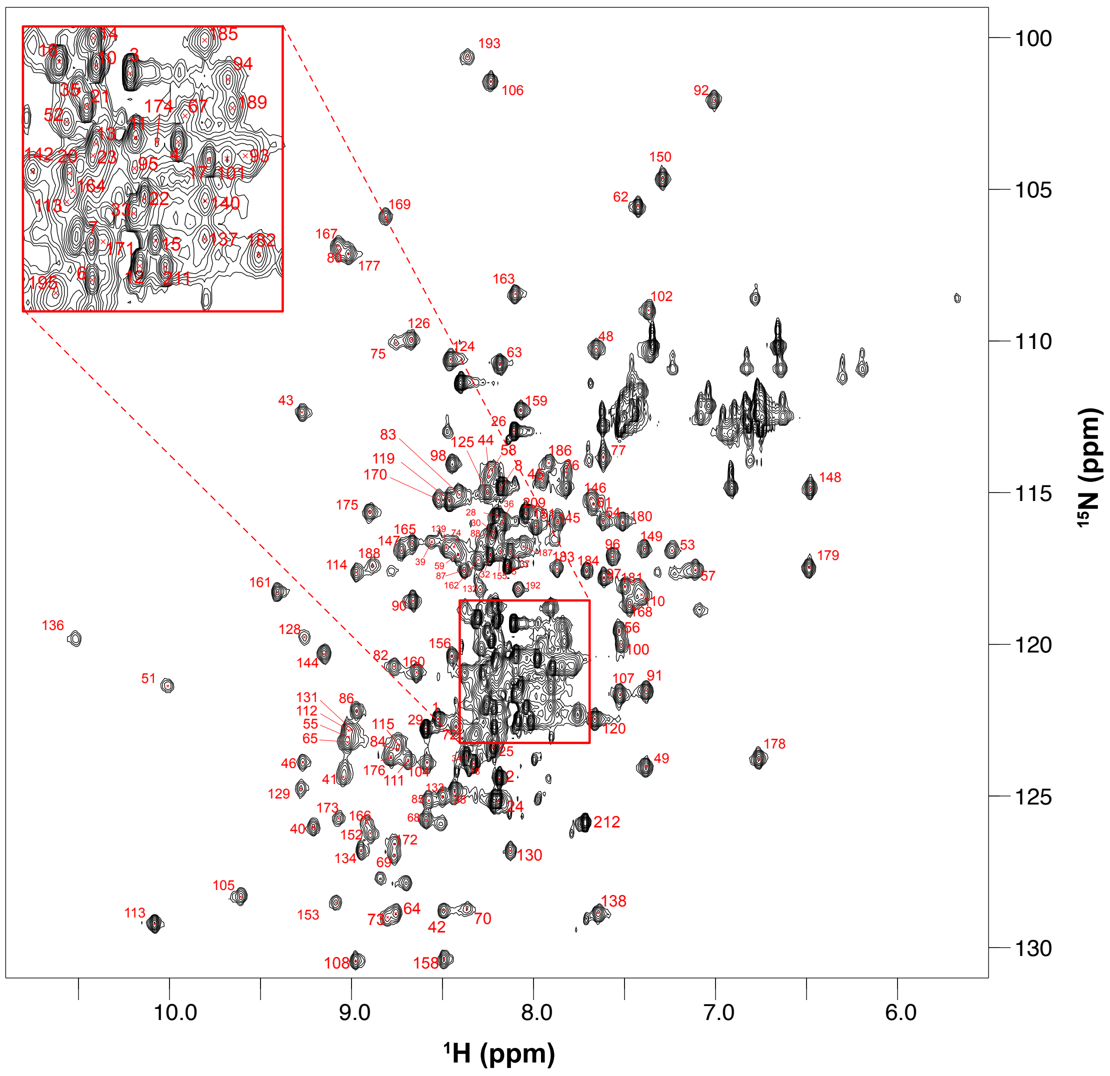


**Supplementary Figure 10.** $^{1}H$-$^{15}N$ HSQC analysis of *Saccharomyces cerevisiae* Pab1 RRM12 with assignments.

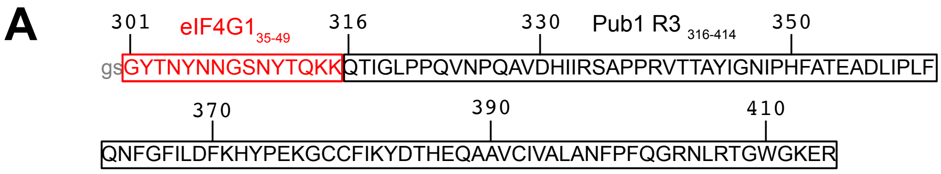


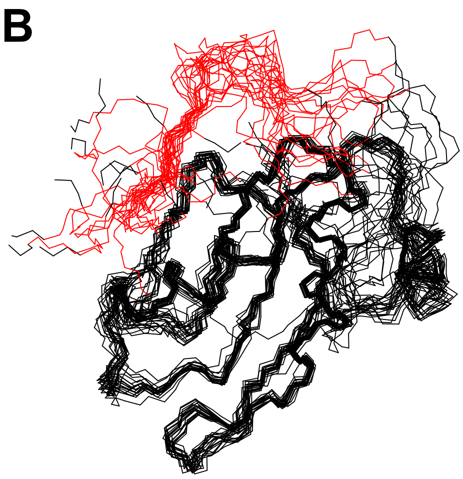


| **Structure calculation statistics** | |
|---|---|
| *NOE-derived NMR restraints* | |
| Intraresidue | 104 |
| Sequential | 62 |
| Medium range (1< i-j < 4) | 46 |
| Long range (i-j > 4) | 0 |
| *Cyana target function* | *0.50 ± 0.06* |
| *Violations* | |
| Distance (> 0.2 Å) (average ± SD) | 9 ± 2 |
| Maximun (Å) | 0.4 |
| *Backbone RMSD (Å)* | 1.39 ± 0.46 |

*Residue range :201-229

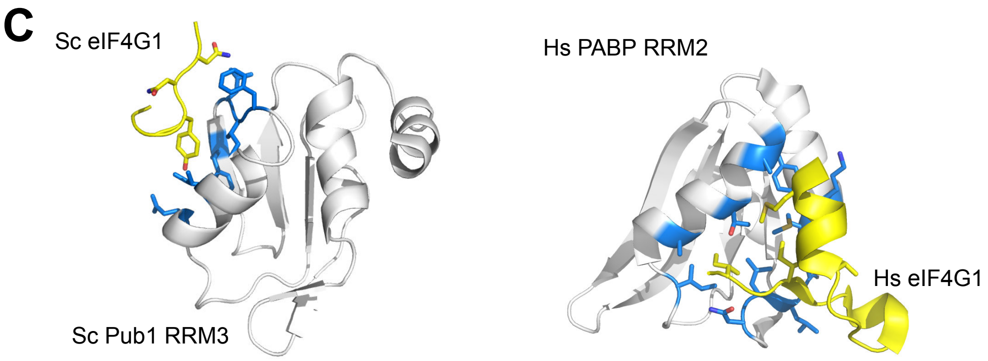


**Supplementary Figure 11. (A)** Sequence of the eIF4G1$_{35\text{-}49}$-Pub1 RRM3 chimera. **(B)** Backbone superposition of the 20 conformers of the NMR structure of the eIF4G1-Pub1 RRM3 chimera (red corresponds to eIF4G1) and the table with structure calculation statistics. **(C)** Comparison of the yeast Pub1 RRM3-eIF4G1 and the human PABP RRM2-eIF4G1 (Safaee et al., 2012) binding modes.
